# Dual impact of PTEN mutation on CSF dynamics and cortical networks via the dysregulation of neural precursors and their interneuron descendants

**DOI:** 10.1101/2023.03.18.533275

**Authors:** Tyrone DeSpenza, Emre Kiziltug, Garrett Allington, Daniel G. Barson, David O’Connor, Stephanie M. Robert, Kedous Y. Mekbib, Pranav Nanda, Ana Greenberg, Amrita Singh, Phan Q. Duy, Francesca Mandino, Shujuan Zhao, Anna Lynn, Benjamin C. Reeves, Arnaud Marlier, Stephanie A. Getz, Carol Nelson-Williams, Hermela Shimelis, Lauren K. Walsh, Jinhui Zhang, Wei Wang, Annaliese OuYang, Hannah Smith, William Butler, Bob S. Carter, Engin Deniz, Evelyn M. R. Lake, Todd Constable, Murat Gûnel, Richard P. Lifton, Seth L. Alper, Sheng Chih Jin, Michael C. Crair, Andres Moreno-De-Luca, Bryan W. Luikart, Kristopher T. Kahle

## Abstract

Expansion of the cerebrospinal fluid (CSF)-filled cerebral ventricles (ventriculomegaly) is the quintessential feature of congenital hydrocephalus (CH) but also seen in autism spectrum disorder (ASD) and several neuropsychiatric diseases. *PTEN* is frequently mutated in ASD; here, we show *PTEN* is a *bona fide* risk gene for the development of ventriculomegaly, including neurosurgically-treated CH. *Pten*-mutant hydrocephalus is associated with aqueductal stenosis due to the hyperproliferation of periventricular *Nkx2.1^+^* neural precursors (NPCs) and CSF hypersecretion from inflammation-dependent choroid plexus hyperplasia. The hydrocephalic *Pten*-mutant cortex exhibits ASD-like network dysfunction due to impaired activity of *Nkx2.1^+^* NPC-derived inhibitory interneurons. *Raptor* deletion or post-natal Everolimus corrects ventriculomegaly, rescues cortical deficits, and increases survival by antagonizing mTORC1-dependent *Nkx2.1^+^* cell pathology. These results implicate a dual impact of PTEN mutation on CSF dynamics and cortical networks via the dysregulation of NPCs and their interneuron descendants. These data identify a non-surgical treatment target for hydrocephalus and have implications for other developmental brain disorders.

**HIGHLIGHTS:**

1. *PTEN de novo* mutations are associated with cerebral ventriculomegaly in autism spectrum disorder (ASD) and congenital hydrocephalus (CH).
2. *Pten*-mutant hydrocephalus is associated with aqueductal stenosis due to the hyperproliferation of medial ganglionic eminence *Nkx2.1^+^* neural precursors and CSF hypersecretion from inflammation-induced choroid plexus hyperplasia.
3. The hydrocephalic *Pten*-mutant cortex exhibits ASD-like network dysfunction due to impaired activity of *Nkx2.1^+^* NPC-derived inhibitory interneurons.
4. mTORC1 inhibition via *Raptor* deletion or early post-natal treatment with rapamycin or everolimus increases survival and ameliorates *Pten*-mutant ventriculomegaly and cortical pathology.

## INTRODUCTION

Da Vinci, echoing Galen 1300 years earlier, maintained that all the brain’s mental functions reside within the cerebrospinal (CSF)-filled ventricular system, as opposed to the brain parenchyma^1,2,3^. Nonetheless, data has only recently emerged that is beginning to shift the commonly held view of the cerebral ventricles from being passive fluid receptacles to more active participants in brain development and function by facilitating multiple cellular, molecular, and biophysical interactions at the brain-CSF interface^4^. For example, the finely-tuned maintenance of intraventricular CSF hydrostatic pressure is essential for morphogenesis of the developing cortical mantle^5^. The precise regulation of neural stem cell growth, proliferation, and fate determination in the fetal ventricular and subventricular zone (V-SVZ), facilitated by the delivery of choroid plexus (ChP)-derived growth factors in the CSF, is critical for neurogenesis, corticogenesis, and synaptogenesis^6,7^. Recently, CSF from juvenile mice has been shown to restore brain function and memory in aged mice by promoting the proliferation of oligodendrocyte progenitors, the largest neural stem cell (NSC) population in the aged brain. Further, the ChP has been shown to secrete signals that extrinsically instruct adult NSC proliferation within the V-SVZ in an age-dependent fashion^8^. These data suggest a better understanding of the “hollow”^2^ of the CSF-ventricular system could help elucidate the development of the adjacent brain parenchyma^9^.

Hydrocephalus (*Gr*., “water on the brain”) is characterized by dilation of the cerebral ventricles (ventriculomegaly). Hydrocephalus often arises from secondary causes such as hemorrhage or infection (acquired hydrocephalus). In the absence of a known antecedent, hydrocephalus is classified as primary, developmental, or congenital hydrocephalus (CH)^10^. Most acquired hydrocephalus cases are attributed to intraventricular CSF accumulation due to a relative mismatch in the ratio of CSF secretion to CSF reabsorption^10,11^. In this context, neurosurgical CSF diversion can normalize ventricular size and improve neurological function. However, these procedures have high rates of technical failure and morbidity^12^ and can fail to improve the associated neurodevelopmental phenotypes of patients, such as intellectual disability, autism spectrum disorder (ASD)^13,14^, and epilepsy^15^. These observations suggest CH is not simply a disorder of impaired fluid homeostasis, but rather a complex developmental brain disorder (DBD) with associated structural and functional deficits of the brain parenchyma. Our incomplete understanding of CH pathobiology and its antiquated anatomically-based taxonomy, while useful for clinical care, hinders improvements in prognostication, treatment stratification, and the development of non-surgical treatments.

Ventriculomegaly is also a common, if often overlooked, associated finding in multiple DBDs and neuropsychiatric disorders, including ASD^16^, idiopathic mental retardation^17^, fragile X syndrome^18^, schizophrenia^19^, and bipolar disorders^20^. ∼20% of children with CH are also diagnosed with ASD – a rate 10-fold higher than the 1.9% rate in the general population^13,21,22^. A retrospective analysis of brain MRI data revealed increased volumes of the cerebral ventricles and of the CSF-producing ChP in ASD patients compared to neurotypical individuals^23^. Ventriculomegaly in these disorders arises during fetal development and can be detected by prenatal ultrasound^17,24,25^. MRI detection of increased extra-axial CSF spaces (“external hydrocephalus”) in neonates can predict a future diagnosis of ASD^26-28^. Thus, ventriculomegaly or enlargement of the other subarachnoid or cisternal CSF spaces could be a useful structural correlate or early radiographic biomarker of aberrant intrinsic cortical function relevant for multiple DBDs and neuropsychiatric conditions. However, the onset, progression, and mechanism of ventriculomegaly in these disorders is poorly understood, in part due to the lack of animal models of ventriculomegaly carrying patient-specific gene mutations.

Recent genomic work has shifted the paradigm of CH pathogenesis from one of “impaired brain plumbing” to one of NSC fate dysregulation in a significant number of cases ^29-31^. Whole-exome sequencing (WES) has shown damaging *de novo* mutations (DNMs) account for ∼20% of sporadic CH patients^29^. Each of the genes in this study harboring a genome-wide significant enrichment of DNMs is a regulator of neural stem cell fate and prenatal neuro-gliogenesis. For example, one of the most mutated genes in CH, *TRIM71*, is an RNA-binding protein in the let-7 pathway that regulates the timing of stem cell progression during early development^4^. CH-associated mutation or neural stem cell-specific *Trim71* depletion in mice causes premature neuronal differentiation at the expense of neural stem cell expansion^32^, thereby depleting the available neural stem cell population for neurogenesis and expansion of the cortical mantle. The resultant cortical hypoplasia leads to a hypercompliant, “floppy” cortex and secondary ventricular enlargement without primary defects in CSF circulation^4^. These data highlight the importance of the precise regulation of periventricular neural stem cells and other precursor cells during fetal development for normal brain biomechanics and CSF homeostasis^30^. Nonetheless, the genetic origins of most sporadic CH cases are unknown, the mechanisms of ventriculomegaly in CH remain obscure, and the function of cortical networks in the context of CH have yet to be studied.

Phosphatase and tensin homolog deleted on chromosome 10 gene (*PTEN*) encodes a dual specificity protein and lipid phosphatase that counteracts the activity of phosphoinositide 3-kinase (PI3K) by dephosphorylating phosphatidylinositol (3,4,5)-triphosphate (PIP3) to phosphatidylinositol (4,5)-bisphosphate (PIP2)^33^, thereby negatively regulating the AKT (protein kinase B)/mechanistic target of rapamycin (mTOR) signaling pathway^34^. This pathway is an essential signaling hub for regulating diverse cellular functions such as growth, metabolism, proliferation, and survival^35,36^. Somatic or germline dysregulation of *PTEN* and other genes in the PI3K/AKT/mTOR pathway have been implicated in a variety of overgrowth syndromes, cancers, and neurological disorders, including ASD, epilepsy, and macrocephaly^37,38^. For example, *PTEN* was recently identified among the top high-confidence ASD risk genes (FDR≤0.1) in an analysis of rare germline *de novo* and case-control variants from ∼12,000 affected ASD probands^38^ corroborating earlier ASD exome studies^39-41^. In mouse models, *Pten* deletion leads to neuronal hypertrophy, hyperexcitability, seizures, and ASD-like behaviors^42-45^. Despite a clear association between *PTEN* loss-mediated mTOR activation and aberrant neuronal morphology and function, the prevalence of *PTEN* mutations in patients with ventriculomegaly or CH has not been comprehensively examined, and the mechanisms of *PTEN*-mutant ventriculomegaly are unknown.

Here, we present a systematic WES analysis that show germline DNMs in *PTEN* are a frequent cause of sporadic, neurosurgically-treated CH. By analyzing the exomes of two other patient cohorts comprising over 8,300 exomes, we found *PTEN* is also commonly mutated in patients with ASD and other NDDs featuring cerebral ventriculomegaly. Using integrative genomics, we found *PTEN* is most highly expressed in the human brain in fetal *Nkx2.1^+^*neural precursors (NPCs) of the subventricular medial ganglionic eminence (MGE) and their post-natal inhibitory interneuron descendants. By engineering mice with a human CH-and ASD-associated *PTEN* mutation, or conditionally deleting *Pten* in embryonic *Nkx2.1^+^* NPCs, we recapitulated the phenotypic continuum of ventriculomegaly exhibited *PTEN*-mutated human patients. Our physiological and cellular experiments results show *Pten*-mutant ventriculomegaly is associated with aqueductal stenosis, or in some cases, complete aqueductal obstruction due to periventricular *Nkx2.1^+^*NPC hyperproliferation, in combination with bumetanide-sensitive CSF hypersecretion due to inflammation-depedent hyperplasia of the ChP epithelium. Notably, fetal genetic inhibition of mTORC1 by *Raptor* knockout or post-natal pharmacologic mTORC1 inhibition by rapamycin or the FDA-approved rapalog Everolimus reduces *Pten*-mutant ventriculomegaly, improves cortical pathology, and increases survival.

Together, these data suggest PTEN is a critical determinant of human CSF homeostasis, ventricular morphogenesis, and cortical function via its regulation of *Nkx2.1^+^* NPCs at the brain-CSF interface, and that this mechanism is likely compromised by PTEN loss-of-function in a significant number of patients with CH and ASD. These results identify a potential non-surgical precision treatment for CH and suggest ventriculomegaly could be a useful radiographic biomarker in other DBDs and neuropsychiatric disorders.

## RESULTS

### Enrichment of *de novo* mutations in PI3K signaling genes in neurosurgically-treated patients with congenital hydrocephalus

To elucidate the genetic determinants of human ventricular size and the cellular and molecular mechanisms of CH, we analyzed WES data from an expanded discovery cohort^29^ of exomes comprising 482 total probands (including 281 proband-parent “trios”) (see Methods). All probands had undergone endoscopic third ventriculostomy or cerebral ventricular CSF shunting for the diversion of CSF (herein defined as “treated CH”) (see **Methods** and **Table S1**). We identified a total of 13 genes with ≥ 2 protein-altering *de novo* variants (DNVs), versus 3.25 genes expected by chance (4.01-fold enrichment; P = 3.64 × 10^−5^ by 1 million permutations; **Table 1**). Greater enrichment of recurrent genes was observed in loss-of-function (LoF)-intolerant genes (pLI > 0.9) with multiple DNVs (241.1-fold enrichment; P = 8.74 × 10^−14^; **Table 1**), supporting these as causal CH disease genes. Five genes, including the PI3K signaling genes *PTEN* and *PIK3CA*, surpassed genome-wide significance thresholds for harboring an excess of protein-damaging DNVs (P-value threshold of 8.6 × 10^−7^ after correction for testing 19,347 RefSeq genes in triplicate by one-tailed Poisson test; **Fig. 1A, and Table 1**). Eight other predictive loss-of-function intolerant genes, including the PI3K signaling gene *MTOR*, also exhibited ≥2 protein-damaging DNVs (**Table 1**).

**Figure 1.**
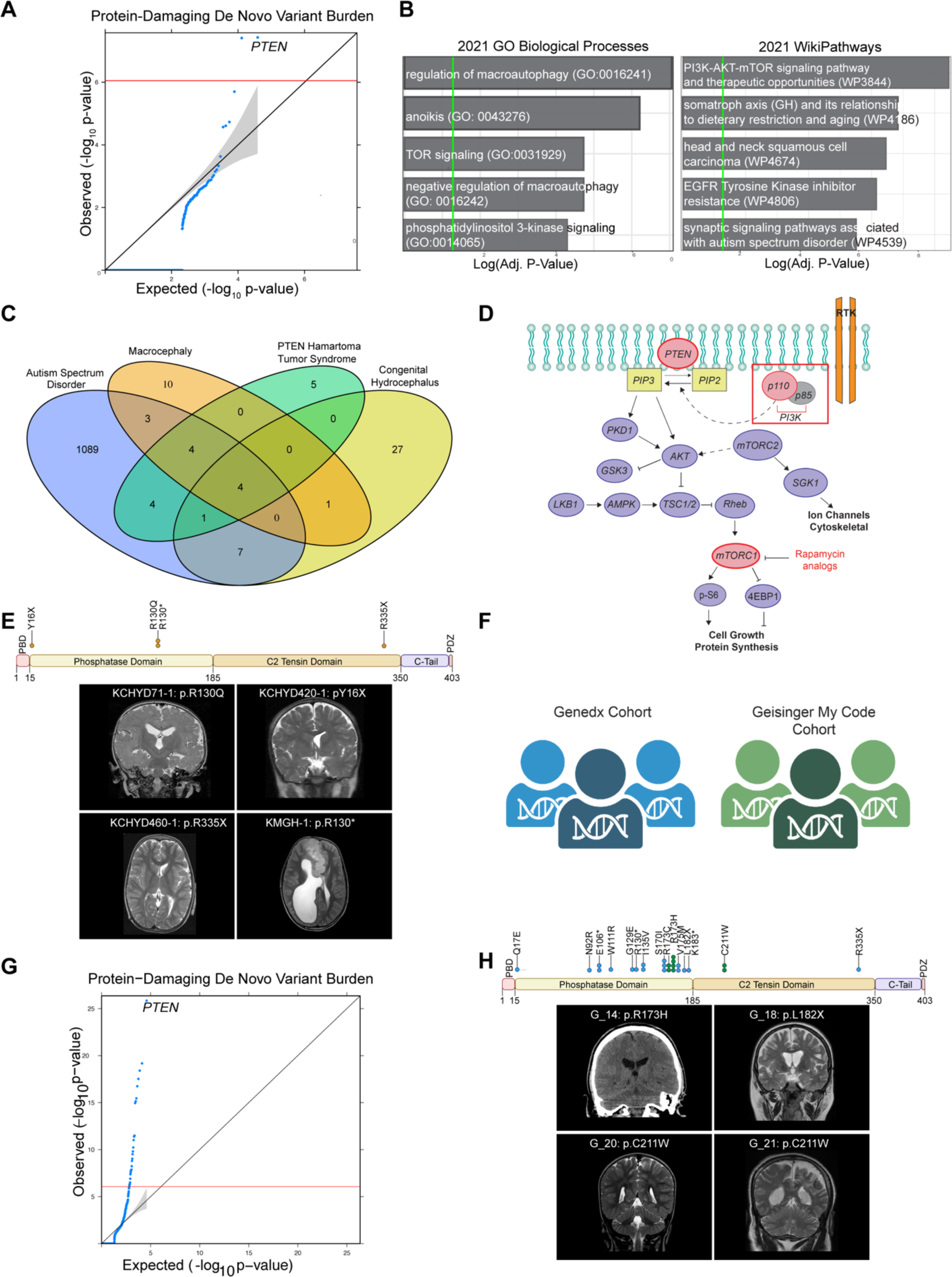
*PTEN* mutation in patients with neurosurgically-treated congenital hydrocephalus, autism, and other neurodevelopmental disorders. (A) Quantile-quantile plot displaying individual variant burdens for each gene identifies genes with a significantly outsized burden. Exome-wide significance, defined as p ≤ 0.05/(19,348*3). (B) Gene Ontology Biological Processes and WikiPathways analyses of possible congenital hydrocephalus risk gene list suggest involvement of the PI3K-AKT-mTOR pathway in congenital hydrocephalus pathogenesis. (C) Possible congenital hydrocephalus gene list significantly overlaps with DisGeNET-curated gene lists related to macroencephaly, autism spectrum disorder, and other neurodevelopmental phenotypes. (D) Schematic of the PI3K-AKT-mTOR pathway with CH-mutated genes highlighted in red. (E) Mapping of all *de novo* variants in *PTEN* identified in the neurosurgically-treated congenital hydrocephalus discovery cohort identifies several recurrent variants which may identify clinically significant residues. Representative radiographic brain imaging of individuals from the neurosurgically-treated congenital hydrocephalus cohort with co-morbid neurodevelopmental disorders (including autism spectrum disorder) and harboring *de novo PTEN* mutations (See table 1). (F) Schematic of GeneDx and Geisinger MyCode Cohorts. (G) Quantile-quantile plot displaying individual variant burdens for each gene identifies genes with a significantly outsized burden in the GeneDx cohort. Exome-wide significance, defined as p ≤ 0.05/(19,348*3).(H) Mapping of all *de novo* variants in *PTEN* identified in the Gene Dx and Geisinger MyCode Cohorts identifies several recurrent variants which may identify clinically significant residues. Representative radiographic brain imaging of individuals harboring damaging PTEN mutations from the Geisinger MyCode Cohort treated ventriculomegaly with co-morbid neurodevelopmental disorders (including autism spectrum disorder; See table 1).

GOrilla and enrichR gene ontology and pathway analysis were performed on those genes with a pLI > 0.9 in ExAC that harbored ≥2 protein-altering DNVs, with at least one protein-damaging DNV (“high-confidence (hc)-CH genes” [9 genes in total]) and ≥1 damaging DNVs (“possible (p)-CH genes” [40 genes in total]) (**Table S1**) (**see Methods**). The most significantly enriched gene ontology (GO) terms were those related to phosphatidylinositol 3-kinase signaling (GO:0014065; 5.21-fold enrichment; modified Fisher’s exact test p = 4.25 × 10^−5^ after Bonferroni correction), the regulation of macroautophagy (GO: GO:0016241; 3.31-fold enrichment; p = 8.86 × 10^−8^), and mTOR signaling (GO: GO:0031929; 7.03-fold enrichment; p = 1.86 × 10^−5^) (**Fig. 1B**). DisGeNET analyses showed genes with DNVs overlapped with those previously implicated in PTEN Hamartoma Tumor Syndrome (Adjusted p = 8.83 x 10^-11^: one-tailed Fisher’s exact test), Macroencephaly (Adjusted p = 3.73 x 10^-10^), and Autistic Disorder (Adjusted p = 3.41 x 10^-7^) (**Fig. 1C**).

10/111 (9.01%) damaging DNVs were identified in genes related to Wikipathway WP3844, “PI3K-AKT-mTOR signaling pathway and therapeutic opportunities” (Adjusted P = 6.58 x 10^-7^, including a total of six LoF and three D-mis DNVs in six genes (**Fig. 1B, Table S1**). Several genes included under the WP3844 term harbored rare, likely pathogenic variants or variants of unknown significance (VUS) in other unrelated CH probands (**Table S1**). Damaging DNVs in genes in the related mTOR signaling term GO:0031929 were enriched in CH cases but not in controls (17.1-fold enrichment; p = 7.72 × 10^−4^) (**Table S1**). Among these, *PIK3CA* and *PTEN* surpassed genome-wide significance thresholds for both DNV enrichment and case-control tests (**Table S1**). mTOR signaling GO:0031929 genes with DNVs mapped to a highly significant genetic and protein-protein interaction network (**Suppl. Fig.1**). These results suggest pathogenic DNVs in *PTEN* and other PI3K signaling genes contribute to a significant percentage of sporadic, treated CH cases.

### *De novo PTEN* mutations in patients with treated hydrocephalus, co-morbid NDDs, and other structural brain defects in a neurosurgery-based cohort

Among the top-ranked hits in our neurosurgically-treated CH cohort, *PTEN* harbored a total of four LoF or damaging DNMs (p.Tyr16X, p.Arg130Gln, p.Arg130*, and p.Arg335X) in unrelated probands ( P = 5.75 × 10^−11^, one-tailed Poisson test for protein-damaging DNVs for expected vs. observed protein-damaging DNVs) (**Fig. 1E, Table S1**). In addition to CSF diversion-dependent hydrocephalus, probands variably exhibited macrocephaly (without megalencephaly), cerebellar tonsillar ectopia, polymicrogyria, and neurodevelopmental delay (**Fig. 1E, Table S1**). For example, KMGH-1 harboring DNM p.Arg130* also had ASD, epilepsy, and a dysplastic corpus callosum (**Fig. 1E, Table S1**). The p.Arg335X CH proband with neurodevelopmental delay had been treated with ventriculoperitoneal CSF shunting, followed by 39 subsequent shunt revision neurosurgeries for hardware malfunction or infection (**Fig. 1E, Table S1**).

DNMs mapped to highly conserved residues the phosphatase domain, C2 Tensin domain of PTEN (**Table S1** and **Fig. 1E**). p.Arg130Gln and p.Arg130* have been previously linked to PTEN hamartoma tumor syndrome (PHTS) (OMIM # 158350), including the Cowden, Bannayan–Riley–Ruvalcaba, and autism-macrocephaly syndrome subtypes^46^. Amino acid residue 130 of PTEN is a site of frequent somatic mutation in cancers of the breast^47^, thyroid^48^, head and neck^49^, and endometrium^50^. p.Arg130Gln, p.Arg130*, and p.Arg335X have been reported in patients with autism-macrocephaly syndrome^51-58^. Notably, these neurosurgically-treated *PTEN*-mutated probands had no previous diagnosis of PHTS, autism-macrocephaly syndrome, or other cancers before WES. These data show rare, damaging germline DNVs in *PTEN* are enriched in neurosurgically-treated CH patients with co-morbid NDDs and other structural brain defects.

### *PTEN* mutations in patients with ventriculomegaly or hydrocephalus and co-morbid ASD and other NDDs in a clinical genetics laboratory cohort and in a healthcare-based cohort

We next assembled a cohort of 2,697 proband-parent trios with cerebral ventriculomegaly, including neurosurgically-treated hydrocephalus, in collaboration with GeneDx, a clinical genetics laboratory comprising over 300,000 exome proband-parent trios^59,60^ (see Methods and **Fig. 1F**). We used DenovolyzeR to assess relative by-gene burden of *de novo* variation. Strikingly, *PTEN* was the top hit among genes harboring enrichment of protein-altering DNMs (P = 3.12 x 10^-30^, xxx) (**Fig. 1G**) far surpassing genome-wide significance thresholds. PTEN contained a total of 17 DNMs, mapped to highly conserved residues of the phosphatase and C2 tensin domains (**Table S1** and **Fig. 1H**). Three of these DNMs (p.His93Arg, p.Pro246Leu, and p.Arg130*) have been previously identified in ASD<u>^61^</u>; the latter is recurrent with unrelated patient KCHYD460-1 in our treated CH cohort. In addition to ventriculomegaly or hydrocephalus, GeneDx *PTEN*-mutant probands variably exhibited macrocephaly, mild to severe neurodevelopmental delay, seizures, ASD, other NDDs, and a variety of other structural brain defects (**Table S1**). These findings replicate those from the treated CH cohort and further corroborate the association of PTEN DNMs with ventriculomegaly and ASD.

We also examined the association of *PTEN* mutation with ventriculomegaly in an outpatient non-surgical cohort using the Geisinger MyCode Community Health Initiative (MyCode), a healthcare-based genomic screening program comprising 169,876 individuals with exome sequencing and linked electronic health records^62^ (see Methods and **Fig. 1H**). Of these, 24 participants had rare, unphased ClinVar pathogenic or likely pathogenic *PTEN* variants mapped to highly conserved residues of the xxx and xxx domains (**Table S1** and **Fig. 1H**), and two recurrent *PTEN* variants (p.Arg130* and p.Arg335*) that were identified in unrelated patients in the treated CH cohort. p.Arg130* was identified in all three cohorts in unrelated patients (**Table S1**). We also queried the Geisinger Autism & Developmental Medicine Institute’s cohort of individuals who consented to the Making Advances Possible (MAP) research protocol. Six additional participants harboring rare, unphased ClinVar pathogenic or likely pathogenic *PTEN* variants identified through routine clinical genetic testing (**Table S1**), mapped to highly conserved residues of the phosphatase and C2 tensin domains (**Table S1** and **Fig. 1E**).

Of these 30 Geisinger patients with *PTEN* variants, 90% had NDDs or neuropsychiatric conditions, including generalized anxiety disorder (50%), depression (37%), intellectual disability/developmental delay (30%), ASD (20%), and/or epilepsy (17%) (**Table S1**). 33% percent of participants had macrocephaly and 30% had a personal history of cancer, most commonly breast (20%). Clinical neuroimaging studies were available for review for 18/30 individuals (60%); among these, 17% had at least mild ventriculomegaly (**Table S1 and Fig. 1H**). Other structural brain defects were noted, including dysgenesis of the corpus callosum, choroid plexus (ChP) hyperplasia, agenesis of the septum pellucidum, and cerebellar tonsillar ectopia. Notably, two unrelated patients with the recurrent *PTEN* p.Cys211Trp variant in the C2 domain each exhibited ventriculomegaly and ChP hyperplasia (**Fig. 1H**), with one proband (G_20) also having co-morbidities of ASD, intellectual disability, and global developmental delay, and the other proband (G_21) having co-morbid epilepsy. These results show rare, damaging *PTEN* germline variants are associated with ventriculomegaly and other structural brain anomalies such as ChP hyperplasia in a non-surgical, healthcare-based cohort that includes patients with ASD and other NDDs.

### *PTEN* is highly expressed in fetal subventricular zone *Nkx2.1^+^* neural progenitors of the medial ganglionic eminence and their post-natal inhibitory interneuron descendants

To gain insight into the cellular and molecular mechanisms by which *PTEN* mutations cause ventriculomegaly, we studied the expression of *PTEN* in the developing human brain using 14 different sc-RNAseq transcriptomic atlases, comprising 144,047 cells from 20 brain regions and 42 cell types across 11 different developmental time points^63^ (**Fig. 2A**). We found that *PTEN* expression was already evident at human post-conception week 4 (PCW4), the time of neurulation, and most highly expressed at PCW 13-16 (**Fig. 2B**, p = 5.22×10^-59^) – an epoch overlapping with peak cortical neurogenesis (PCWs 7-24)^64^. Further examination of the PCW 13-16 epoch revealed highest expression of *PTEN* in the medial ganglionic eminence (MGE) (**Fig. 2C**), an embryonic neural progenitor domain within the ventral telencephalon abutting the ventricular system that gives rise to most GABAergic interneurons in the cerebral cortex^65,66^.

**Figure 2.**
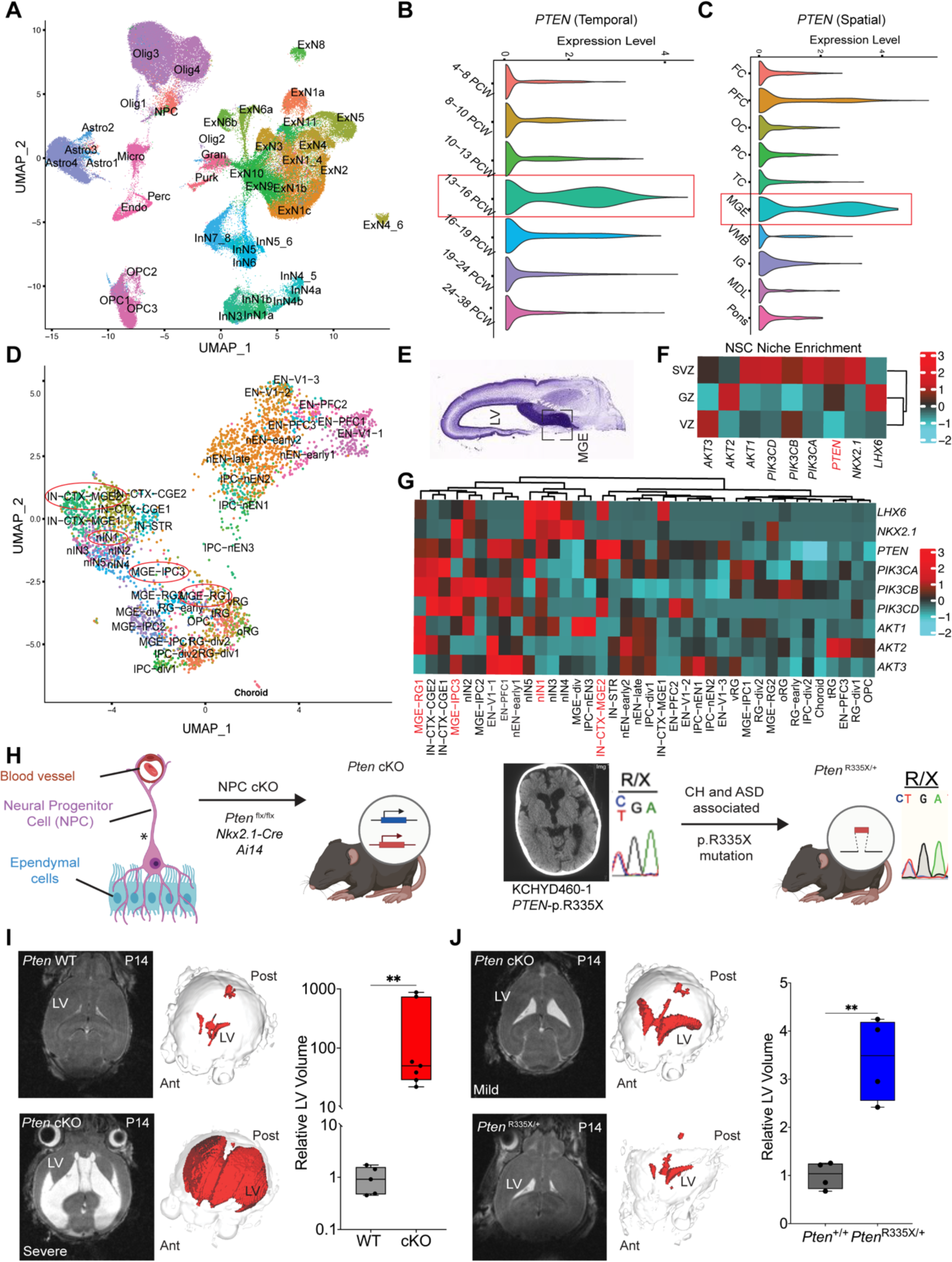
Fetal *Nkx2.1^+^* neuroprogenitors are a critical spatio-temporal locus of pathobiology in PTEN-mutant ventriculomegaly. (A) UMAP clustering of human fetal brain cells using <u>S</u>patio-<u>T</u>emporal cell <u>A</u>tlas of the human <u>B</u>rain (STAB), colored by cell type. (B)-(C) Violin plots of *PTEN* gene expression in the human brain using <u>STAB</u>. PTEN was most highly expressed during PCW 13-16 (Avg Log_2_FC=1.64, p=2.17e-63). Within that timeframe, PTEN was mostly highly expressed in the Medial Ganglionic Eminence (MGE). (D) UMAP clustering of developmental human brain neuro-progenitor cells using a publicly available spatiotemporal dataset (Nowakowski et al.), colored by cell type. (E) Fetal brain image from the human BrainSpan atlas to illustrate the proximity of the MGE to the lateral ventricle (LV). (F) Gene expression heatmap of *NKX2.1*, *LHX6, PTEN* and *PI3K* signaling genes in the subventricular zone (SVZ), ventricular zone (VZ) and germinal zone (GZ), showing enriched expression of these genes in SVZ. (G) Gene expression heatmap of *NKX2.1*, *LHX6*, PTEN, and *PI3K* signaling genes across different cell types, showing highest PTEN expression in MGE-derived cortical inhibitory neurons (IN-CTX2-MGE), Striatal Neurons (IN-STR), MGE progenitor cells (MGE-IPC3) and MGE newborn neurons (nIN1). (H) Identification of the *de novo PTEN*-p.R335X mutation in hydrocephalic proband KCHYD460-1 with history of 50+ ventriculoperitoneal shunts motivated generation of a humanized *Pten*^R335X/+^ mouse model by CRISPR-Cas9 (<u>Methods</u>). (I) Additional *Pten* cKO mouse model generated by conditional deletion of *Pten* in *Nkx2.1*-expressing neural stem cells via *Nkx2.1-Cre* mediated recombination of Pten^flx/flx^ alleles with the Ai14 tdTomato reporter of *Nkx2.1-Cre* expression (<u>Methods</u>). (J) Representative Brain MRI and associated 3D reconstructions of the ventricular system demonstrates ventriculomegaly in *Pten*^R335X/+^ and *Pten* cKO mutant mice relative to littermate controls (*Pten* WT vs. *Pten* cKO mice from P10-P14; *Pten^+/+^* vs. *Pten*^R335X/+^ mice at P14). (K) Quantitation of ventricular volume from MRI scans of *Pten* mutant mice vs. controls at P10-P14. Significance was tested by two-sided, unpaired *t*-test. Early and Late Born Excitatory Neuron PFC (EN-PFC), Early and Late Born Excitatory Neuron V1 (EN-V1), CGE/LGE-derived Inhibitory Neurons (IN-CTX-CGE), MGE-derived Ctx Inhibitory Neuron (IN-CTX-MGE), Striatal Neurons (IN-STR), Dividing Intermediate Progenitor Cells RG-like (IPC-div), Intermediate Progenitor Cells EN-like (IPC-nEN), Dividing MGE Progenitors (MGE-div), MGE Progenitors (MGE-IPC), MGE Radial Glia (MGE-RG), Mural/Pericyte (Mural), Newborn Excitatory Neuron - Early Born (nEN-early), Newborn Excitatory Neuron - Late Born (nEN-late), MGE newborn neurons (nIN), Oligodendrocyte Progenitor Cell (OPC), Outer Radial Glia (oRG), Dividing Radial Glia (RG-div), earlyRG (RG-early), Truncated Radial Glia (tRG), Unknown (U), Ventricular Radial Glia (vRG). Lateral Ventricle (LV), Anterior (Ant), Posterior (Post).

To study *PTEN* expression at earlier human developmental points in neural progenitor cells, we used a comprehensive spatiotemporal sc-RNAseq human atlas comprising 4261 cells from 3 laminar zones and 48 different cell types spanning PCWs 5-20^67^ (**Fig. 2D**). We found the highest expression of *PTEN* in the subventricular zone (SVZ) of the MGE, a proliferative region situated at the lateral wall of each lateral ventricle containing NPCs, which divide to produce neurons in the process of neurogenesis^66^ (**Fig. 2E**). In the SVZ, PTEN was most highly expressed in *Nkx2.1*-positive MGE neural precursor cells (Avg Log_2_FC = 0.53), MGE newborn neurons (Avg Log_2_FC = 0.5), and MGE-derived cortical inhibitory neural progenitors (Avg Log_2_FC = 1.05) (**Fig. 2F-G**). Nkx2.1 is required for normal patterning of the MGE and for the specification of the parvalbumin- and somatostatin-expressing cortical interneurons^66^. These results show that human *PTEN* is most highly expressed in fetal *Nkx2.1+* NPCs of the MGE SVZ and their post-natal inhibitory interneuron descendants.

### ASD- and CH-associated *Pten* mutation or *Pten* deletion in fetal *Nkx2.1^+^* neural progenitors cause murine ventriculomegaly

First-trimester NPCs, rather than components of the CSF secretory-transit-reabsorption machinery *per se*, have been identified as an unexpected spatio-temporal locus of CH disease gene convergence in the human brain, and as a promising cell type for study of CH-associated gene mutations^29,31^. Fetal *Nkx2.1^+^* neural progenitors within the MGE, located adjacent to the CSF-containing cerebral ventricles, give rise to ∼70% of GABAergic interneurons in the mature cerebral neocortex^68^,^66^ and have been implicated in the proposed dysregulated excitatory/inhibitory (E/I) tone characteristic of ASD^69^, including in *PTEN*-mutated ASD patients. We hypothesized PTEN loss-of-function in *Nkx2.1*^+^ NPCs drives both the anatomical (ventriculomegaly) and functional pathology (ASD and other NDDs) of *PTEN*-mutated patients.

To test this hypothesis, we conditionally deleted *Pten* in C57BL6/j mouse *Nkx2.1*^+^ NPCs using Cre-lox methodology (see Methods). We also studied the fate-mapped expression of *Pten* using a tdTomato reporter (*Nkx2.1*-Cre; *Pten^flx/flx^*; *Ai14^flx/+^*, herein referred to as “*Pten* conditional knockouts [*Pten* cKOs]”) (**Fig. 2H,** and **Methods**). The *Nkx2.1*-Cre BAC transgenic drives expression, beginning approximately embryonic day 10.5 (E10.5), in the VZ/SVZ of the MGE and POA^70^. Consistent with this, we noted robust *Nkx2.1*-Cre-dependent tdTomato expression and efficient Pten protein depletion in early NPCs of the VZ/SVZ and their progeny in *Nkx2.1*^+^-Cre lineage domains of the MGE and POA by E12.5 (**Suppl. Fig 2, A-B**). Live-brain MRI showed significant ventriculomegaly in *Pten* cKO mice already by P1, the earliest time point examined (**Suppl. Fig 3**). Quantitative brain MRI showed progressive macrocephaly and hydrocephalus ranging from mild to severe in 100% of *Pten* cKO mice by P14. All *Pten* cKO mice exhibited unprovoked, spontaneous seizures starting ∼P17 (**Suppl. Video 1**) and died around the time of weaning (∼P21).

**Figure 3.**
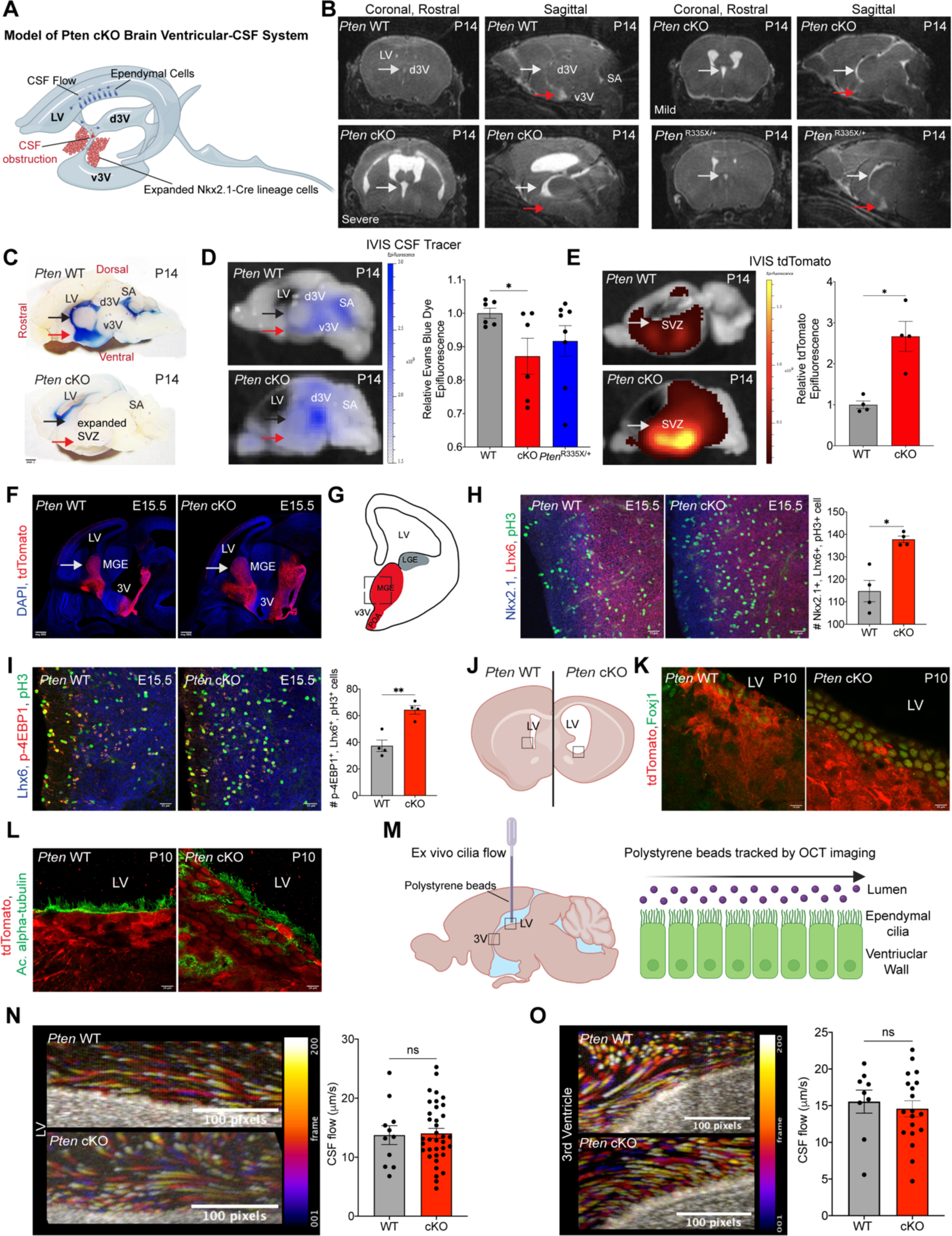
*PTEN*-mutant ventriculomegaly results from aqueductal stenosis due to hyperproliferation of Nkx2.1^+^ neuroprogenitors. (A) Schematic modeling obstructed CSF transport in *Pten* cKO mice.(B) Representative Brain MRI at P14 reveal increased size of lateral ventricles (LV) and dorsal third ventricle (d3V) of the ventricular systems in *Pten*^R335X/+^ and *Pten* cKO mutant mice. White arrows highlight increased d3V caliber in *Pten* mutant mice. Red arrows highlight decreased ventral third ventricle (v3v) caliber in *Pten* cKO mice. (C) Representative midsagittal images of dissected P14 brains from *Pten* WT and *Pten* cKO mice ten minutes post-ICV injection of CSF tracer Evan’s Blue(1%). Black arrows highlight decreased CSF tracer in d3V of *Pten* cKO mice. Red arrows highlight decreased CSF tracer in v3V of *Pten* cKO mice. (D) Representative images of ICV-infused CSF tracer with quantitation of pseudo-color epifluorescence at P14 reveal decreased CSF tracer in the v3V of *Pten* cKO mice. (E) Images and quantitation of pseudo-color epifluorescence superimposed on white images of midsagittal brain dissections from *Pten* WT and *Pten* cKO mice reveal significantly increased endogenous *Nkx2.1-Cre*-dependent tdTomato expression (red) in cells of SVZ and ventral brain. (F) E15.5 midsagittal brain sections from *Pten* WT and *Pten* cKO mouse embryos reveal *Nkx2.1-Cre-*dependent endogenous tdTomato expression (red) in periventricular MGE cells along ventral regions of developing lateral and third ventricles. White arrows highlight increased MGE size in *Pten* cKO. (G) Schematic of E15.5 mouse brain with lateral ventricle (LV) close to lateral ganglionic eminence (LGE) and to medial ganglionic eminence (MGE). (H)–(I) Immunofluorescent labeling and quantitation of nkx2.1 (NSC marker), lhx6 (neuronal-specification NPC marker), p-H3 (marker of cell mitosis), p-4EBP1 (mTor pathway activation marker) in ventral MGE of coronal brain sections from E15.5 *Pten* WT and *Pten* cKO mice. Two-sided, unpaired *t*-test. (J) Schematic of coronal mouse brain section, denoting the location of Nkx2.1-lineage ependymal cells in the ventral domain of the LV. (K) Immunofluorescent imaging of coronal brain section labeled with tdTomato (red), Foxj1 (green) shows differentiated Nkx2.1-lineage ependymal cells in *Pten* cKO and WT littermate controls at P10. (L) Immunofluorescent imaging of coronal brain section labeled with tdTomato (red), Acetylated alpha-tubulin (green) reveals matured ependymal cilia in P*ten* cKO mice at P10. Scale bars, 100 µm. (M) Schematic of OCT measurement of CSF flow speeds of polystyrene beads placed upon ventricular explants. (F)-(G) Representative flow polarity maps demonstrate time-lapse bead trajectories (by temporal color coding) in ventricular brain explants from P*ten* cKO and litter mate control WT mice at P10 with quantitation of local CSF flow speeds at ventricular walls of the lateral ventricle and 3^rd^ ventricle. Color bar represents color versus the corresponding frame in the color-coded image. Scale bars, 100 pixels. Error bars, mean ± sem; each symbol represents one animal. *p<0.05, **p<0.01, ***p<0.001, ****p<0.0001, ns = not significant.

Similarly, C57BL6/j mice engineered with the orthologous *PTEN* Arg335* mutation (*Pten^R335*/R335*^*) previously described in ASD^55^ and also detected in one of our treated CH patients (KCHYD460-1) with co-morbid developmental delay who had undergone > 39 shunt revisions, failed to survive beyond E13.5 (data not shown). Although *Pten^R335*/WT^* mice survived into adulthood, live-brain MRI showed >50% of mice exhibited at least mild ventriculomegaly at P14 (**Fig 2J**). Similar to *Pten* cKO mice, exhibited spontaneous, unprovoked seizures in adulthood (**Suppl. Video 2**). These data show that an ASD- and CH-associated *Pten* mutation or *Pten* deletion in fetal Nkx2.1+ NPCs cause murine ventriculomegaly or hydrocephalus, recapitulating the brain anatomy of human patients.

### *Pten*-mutant hydrocephalus is associated with aqueductal stenosis due to hyperproliferation of ***Nkx2.1^+^* neural progenitors of the subventricular zone MGE**

To begin to elucidate the mechanism of *PTEN*-mutant hydrocephalus, we studied CSF dynamics in *Pten* mutant mice. Hydrocephalus can be categorized as non-obstructive or obstructive, depending on communication or non-communication, respectively, of CSF flow through the Sylvian aqueduct connecting the 3rd and 4th ventricles^10^ (**Fig. 3A)**. The orthologous structure in mice is situated at the more ventrally-positioned 3rd ventricle adjacent to the MGE and POA, with connections to a dorsal 3rd ventricle and Sylvian aqueduct^71^ (**Fig. 3A**). High-resolution imaging and *in vivo* MRI showed *Pten* cKO mice at P14 consistently exhibited either stenosis (narrowing) or complete obstruction at the ventral 3rd ventricle (v3V), with an increased SVZ, as documented (**Fig. 3B, 3C)** and quantified with In Vivo Imaging System (IVIS) imaging of dissected P14 brains intraventricularly infused with the CSF tracer dye, Evans Blue (**Fig. 3C, 3D**). In contrast, *Pten^R335*/WT^* mice with ventriculomegaly at P14 largely had patent v3Vs **(Suppl. Fig 4**).

**Figure 4.**
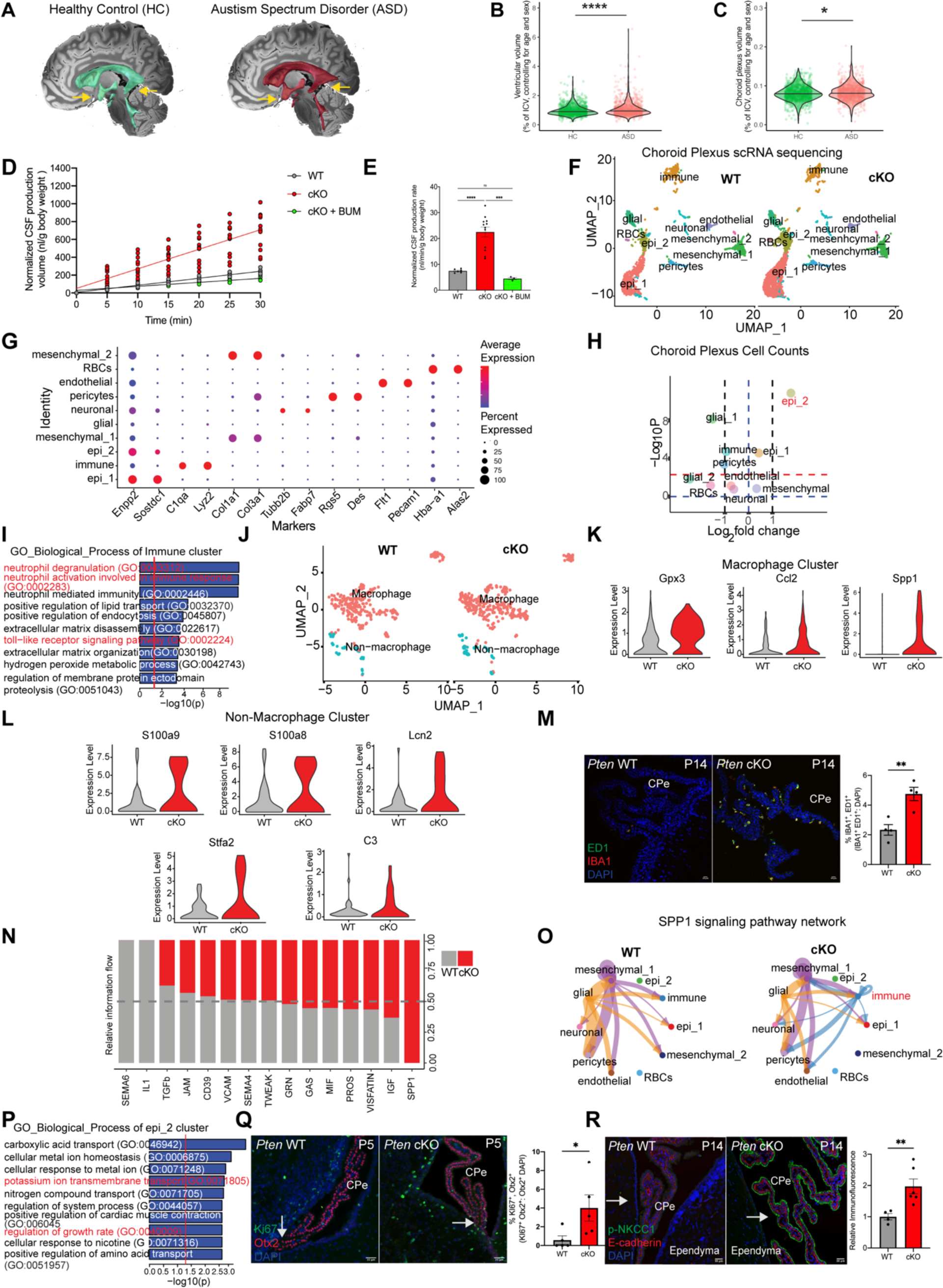
*PTEN*-mutant mice have increased CSF production from an inflamed and hyperplastic choroid plexus. (A)-(C) Representative 3D MRI reconstructions and quantitation of intracranial volume-normalized ventricular volume and choroid plexus volume from 920 patients with autism (ASD) and 1003 healthy controls (HC) from the Autism Brain Imaging Data Exchange (ABIDE) collections, controlled for age and sex. Two-sided, unpaired *t*-test. (D)-(E) Body-weight normalized rate of CSF production in *Pten* WT, *Pten* cKO, and Pten cKO mice treated with bumetanide (BUM). One way Anova. (F) UMAP clustering of ChP cells from *Pten* WT and *Pten* cKO mice, colored by cell type. (G) Expression of cell type markers by scRNA-seq of the ChP. The color indicates average expression level in expressing cells, the size of the circles indicates the proportion of cells expressing a particular marker, which is indicated by the columns, for each cell type cluster rows. (columns) (color) and proportion of expressing cells (circle size) across marker genes (columns). (H) Volcano plots depicting differences of cluster abundance in *Pten* cKO ChP cells compared to control, plotting fold change (log_10_) against *p* value (−log_10_) based on beta-binomial regression. Red horizontal dashed line indicating significance threshold reveals increased cells within “epi_2” subcluster of *Pten* cKO mice. (I) GO biological processes pathway analysis for most highly upregulated genes in *Pten* cKO immune cells vs. WT immune cells. Dotted red line indicates threshold p-value of 0.05. (J) Immunofluorescent labeling and quantitation of co-labeled IBA1 (microglia/macrophage marker) and CD68 clone ED1 (macrophage/monocyte marker) in choroid plexus epithelium of *Pten* WT and *Pten* cKO mice. (K) Further clustering of the immune cells cluster revealed macrophage and non-macrophage cells. (L) Violin plots demonstrating expression of S100a8 (Avg Log_2_FC=2.32, P = 9.80e-4), S100a9 (Avg Log_2_FC=2.10, P = 2.15e-3), Lcn2 (Avg Log_2_FC=2.62, P = 1.18e-4), Stfa2 (Avg Log_2_FC=4.25, P = 1.53e-4), C3 (Avg Log_2_FC=0.43, P = 5.02e-4), in non-macrophage immune cell clusters comparing PTEN cKO condition to control. (M) Violin plots demonstrating expression of Gpx3 (Avg Log_2_FC=0.61, P = 1.05e-12), Ccl2 (Avg Log_2_FC=0.68, P = 1.1e-3), and Spp1 (Avg Log_2_FC=0.91, P = 2.27e-7) in macrophage cell clusters comparing *Pten* cKO cells to control.(N) Significant signaling pathways ranked based on differences in the overall information flow within the inferred networks between epithelial and immune cells, comparing Pten cKO cells to control. The overall information flow of a signaling network is calculated by summarizing all communication probabilities in that network. Rows with high red-to-gray ratio indicate higher ligand-receptor pathway activity for *Pten* cKO cells. (O) Circle plot comparing SPP1 pathway ligand-receptor interactions among cell groups from Pten WT and Pten cKO mice. Edge width represents communication strength. (P) GO biological processes pathway analysis for top gene markers of epithelial sub-cluster epi_2. Dotted red line indicates threshold p-value of 0.05. (Q) Immunofluorescent labeling and quantitation of Ki67 (proliferation marker), Otx2 (choroid plexus epithelial cell marker) in *Pten* WT and *Pten* cKO mice. Two-sided, unpaired *t*-test. (R) Immunofluorescent labeling of p-NKCC1 and E-cadherin in choroid plexus epithelium of P14 *Pten* WT and *Pten* cKO mice, with quantitation of p-NKCC1 fluorescence relative to *Pten* WT control. Two-sided, unpaired *t*-test. Error bars, mean ± sem; each symbol represents one animal. *p<0.05, **p<0.01, ***p<0.001, ****p<0.0001, ns = not significant.

IVIS imaging in *Pten* cKO mice also revealed significantly increased tdTomato reporter fluorescence in *Pten* cKO mice surrounding the lateral ventricle and v3V, suggesting accumulation of *Pten*-deleted cells in the VZ/SVZ and ventral brain, vestiges of the embryonic MGE and POA (**Fig. 3E**). PI3K pathway genes regulate cell growth and proliferation in multiple tissues^72^, including VZ/SVZ NPCs^73,74^. Germline or mosaic *PTEN* mutations have been identified in multiple brain and body overgrowth syndromes that also predispose to cancer^75^, and somatic *PIK3CA* or *MTOR* GoF mutations and *PTEN* LoF mutations drive tumorigenesis by increasing mTOR signaling^76^. mTORC1 promotes NPCs proliferation and differentiation via phosphorylation of Eukaryotic initiation factor 4E-binding proteins (4E-BPs)^77,78^. Further, *Mtor* deletion in *Nkx2.1^+^*-Cre-expressing embryonic NPCs decreases cell proliferation and MGE size at E15.5^79^. We hypothesized that *Pten* deletion in *Nkx2.1^+^* NPCs causes an mTORC1-dependent increase in NPC proliferation and an associated expansion of the periventricular MGE.

To test this hypothesis, we performed quantitative immunofluorescence confocal microscopy of sectioned *Pten* cKO mouse brains and their wild-type littermate brains at E15.5, the time of highest *PTEN* expression in the PCW 13-16 human brain^80^, using antibodies directed against p-H3 (a mitotic index marker) and Lhx6 (a marker of differentiating NPCs). We found the signal of both p-H3 and Lhx6 were significantly upregulated in *Nkx2.1^+^* NPCs of the ventral MGE in *Pten* cKO mice versus controls (**Fig. 3F,3H**). *Pten* cKO NPCs also exhibited increased co-immunostaining of p-H3 and p-4E-BP, indicative of mTORC1 activation and cell hyperproliferation (**Fig. 3I**). Together, these results suggest that PTEN LoF results in mTORC1-dependent hyperproliferation of *Nkx2.1^+^*NPCs that leads to expansion of the ventral MGE, associated compression and stenosis or obliteration of the aqueduct, and a consequent reduction or complete block of CSF flow that leads to obstructive hydrocephalus.

### *Pten*-mutant ventriculomegaly is associated with CSF hypersecretion due to inflammation-associated hyperplasia of the ChP epithelium

Some *PTEN*-mutant CH patients lack complete aqueductal blockage (see **Fig.1H, Table S1**), similar to findings in *Pten* cKO and *Pten*^R335X/+^ mice (see Fig. 3B, 3D). Other *PTEN*-mutant patients with ASD or other NDDs have communicating ventriculomegaly and do not require CSF diversion to survive^81^. An increased frequency of ventriculomegaly has been reported in ASD patients^23,82-85^, including patients with *PTEN*-mutant ASD and macrocephaly^86^. In a cohort of 920 patients with ASD and 1003 age- and sex-matched controls from the Autism Brain Imaging Data Exchange (ABIDE) collection^87,88^, we first corroborated the association of ventriculomegaly with ASD using the convolutional neural network SynthSeg+ to automatically segment subjects’ brain MRIs^89^. We found that ventricular volumes were significantly larger in ASD than in controls (F=32.4, p=1.5e-8; **Fig. 4A, 4B**). FastSurferCNN analysis of the same cohort^90^ showed choroid plexus (ChP) volumes were significantly larger in ASD than in controls (F=5.7, p=0.017; **Fig. 4A,4C**), as previously reported^23,91^. These data implicate mechanisms other than, or in addition to, obstructive aqueductal stenosis in CH- and ASD-associated *PTEN*-mutant ventriculomegaly, including dysregulation of the ChP.

Ventriculomegaly can also result from defective motile-cilia function on the ependymal surface^92,93^ or from increased CSF secretion by the ChP epithelium^94,95^. To examine ependymal motile ciliary function in our *Pten* model, we adopted the optical coherence tomography (OCT) imaging approach we previously validated in *Xenopus*^96^ and mice^4^. Ependymal flow dynamics are visualized and quantified in these experiments by placing polystyrene beads on live preparations of ventricular explants containing the LV and 3V (**fig. S3, F-G**). *Ex vivo* ciliary beating by ependymal cells produces currents that drive bead movement, which can be tracked quantitatively by OCT imaging and calculated as a proxy for microfluidic CSF flow^96^. Application of ciliobrevin, an inhibitor of the dynein motor protein that powers the beating of motile cilia^97^, reduced ependymal flow in the lateral ventricle, and toxin washout restored bead movement, confirming ciliary motion as required for ependymal flow in this model (data not shown).

We then examined ependymal CSF flow in the enlarged LVs and d3Vs of *Pten* WT and cKO mice by OCT imaging at P10. We found cilia-generated CSF flow was unaffected in hydrocephalic *Pten* cKO mice compared to WT controls (**Fig. 3M-0**). Moreover, confocal immunofluorescent microscopic detection of the axonemal protein of matured motile cilia, acetylated α-Tubulin ^98^, and of the ependymal cell differentiation and ciliogenesis marker, Foxj1^98^, revealed morphologically normal motile cilia of differentiated ependymal cells in cKO *Pten* littermates at P10, the timepoint at which *Pten* cKO mice have profound ventriculomegaly and development of multiciliated ependymal cells is complete^99,100^ (**Fig. 3J-L**). Additionally, *Pten* cKO mice displayed obstructive hydrocephalus at P1 (**Suppl. Fig 3**) before ependymal cilia achieve significant functionality within the ventricular system at ∼P3-P10^101^. Together, these data suggest that defects in ependymal cilia-generated CSF flow are not a major driver of *Pten*-mutant ventriculomegaly.

To assess the contribution of increased CSF production to development of hydrocephalus in *Pten*-mutant mice, we directly measured CSF secretion rate in live animals. This validated method^102,103^ applies mineral oil block of CSF exit from the third ventricle at the level of the Sylvian aqueduct, preventing contributions to measured CSF production from CSF reabsorption pathways distal to the block^102,103^, and thereby allowing measurement of *bona fide* lateral ventricle CSF secretion by the ChP^103^. Strikingly, results showed *Pten* cKO mice exhibited a ∼3-fold increase in CSF secretion relative to controls (**Fig. 4D, 4E**). ICV-delivered bumetanide, an inhibitor of ChP-mediated CSF secretion^103-105^, reduced CSF hypersecretion in *Pten* cKO by >70% (**Fig.4E**). These results suggest that non-obstructive ventriculomegaly in *Pten* cKO mice is driven by an increase in bumetanide-sensitive ChP-mediated CSF secretion, which may also potentially exacerbate obstructive Pten-mutant ventriculomegaly.

### *PTEN*-mutant CSF hypersecretion is associated with inflammation-associated ChP hyperplasia and increased Spak-Nkcc1 phospho-activation

ChP-mediated CSF hypersecretion can occur in the context of ChP hyperplasia^95,106-108^, ChP papilloma or carcinoma^109-111^, and ChP inflammation^112,113^. Notably, the frequency of ChP hyperplasia (see **Fig. 4A-C** above) and ChP inflammation is increased in ASD patients^23,82-85^, including those with *PTEN*-mutant autism, macrocephaly, and acetazolamide-sensitive intracranial hypertension^86^. *Nkx2.1^+^*-Cre-dependent tdTomato reporter expression was not detected in the ChP of *Pten* cKO mice (**Suppl. Fig. 5**). Consistent with this observation, Pten immunostaining was detected in the ChP of both WT and *Pten* cKO mice (**Suppl. Fig. 5**). These data suggest a non-cell autonomous impact on ChP secretory function in the mechanism of PTEN-mutant ventriculomegaly.

To gain insight into this mechanism, we first performed single-cell RNA sequencing (scRNAseq) on microsurgically-dissected ChP from *Pten* WT (control) and *Pten* cKO mice (**Fig. 4F**). We then compared the immune and epithelial cell expression profiles using established gene markers of specific cell subclusters (**Fig. 4F, 4G**)^114,115^. The immune cell clusters consisted of 274 control and 345 *Pten* cKO ChP immune cells. The epithelial cell clusters consisted of 864 control and 1901 *Pten* cKO ChP non-immune cells. Clustering of the cell populations in control ChP yielded findings resembling those previously reported^115^.

Compositions of the ChP immune cell sub-clusters were similar in the ChP from *Pten* WT and cKO mice (**Fig. 4H**). However, GO analyses of the most differentially expressed genes in the macrophage and non-macrophage sub-clusters revealed significant enrichment in *Pten* cKO ChP immune cells of pathways related to innate immune cell activation, including neutrophil activation, chemotaxis, and toll-like receptor signaling, (**Fig. 4I**). Among the most highly up-regulated genes in the *Pten* cKO ChP macrophage subcluster were cytokines and other regulators of innate immunity encoded by *Ccl2*, *Spp1*, and *Gpx3* (**Fig. 4J, 4K**). Similarly, *S100a8*, *S100a9*, *Lcn2*, *Stfa2*, and *C3* were among the most up-regulated genes in the non-macrophage subcluster relative to controls (**Fig. 4J, 4L**).

In line with these -omics findings, *Pten*-mutant mice exhibited an accumulation of CD68-, clone ED1-, and Iba1-co-positive immune cells indicative of activated macrophages at the apical, CSF-facing ChP surface, as well as in the ChP stromal compartment between the basolateral membrane and endothelium (**Fig. 4M**). Iba1^+^ cells likely include immune cells recruited from the periphery^115,116^ and CNS border-associated macrophages (BAMs), consisting of epiplexal (Kolmer) and stromal subtypes. In contrast to their quiescent appearance in control animals^115^, BAMs in *Pten* cKO mice exhibited amoeboid-like shape, increased circularity, and increased immunopositivity for CD68 and clone ED1^117^ (**Fig. 4M**), all indicative of increased phagocytic activity^118^. These results demonstrate robust ChP inflammation in the context of *Pten*-mutant hydrocephalus.

Osteopontin (*Spp1*) and IGF ligand-receptor pathways were the most significantly activated pathways linking ChP immune cell and epithelial clusters and in *Pten c*KO mice (**Fig. 4N**), as revealed by CellChat^119^, a scRNAseq tool used for inferring intercellular communication networks based on the expression of known ligand-receptor pairs in different cell clusters. Osteopontin, a Th1 cytokine, through interactions with integrins^120^ and activation of PI3K-Akt-mTOR signaling^121,122^, promotes immune cell migration, adhesion, and survival in chronic inflammation and autoimmune disease^123,124^ and metastasis^125^. Spp1 signals were sent mainly from ChP immune cells to ChP epithelial cells (**Fig. 4N, 4O**). Cell-cell communication analysis revealed ligand-receptor pair Spp1-Cd44 among the main ligand-receptors contributing to the Spp1 communication pathway (**Fig. 4N, 4O**). These results implicate signaling from ChP immune cells to ChP epithelial cells in the mechanism of ChP CSF hypersecretion in *Pten*-mutant hydrocephalus.

Inflammation drives epithelial cell proliferation and carcinogenesis^126^ and also stimulates the vectorial transport of ions and fluid across epithelia epithelia ^127-129^, including the the ChP epithelium^103^. Comparison of ChP-cell sub-clusters from *Pten* WT and *Pten* cKO mice revealed a significant increase in the “epi_2” epithelial sub-cluster (**Fig. 4H**, volcano plot, Log_2_FC=0.80, p=1.51e-10), indicative of ChP epithelial cell hyperplasia. GO analyses of the most differentially expressed genes in the epi_2 sub-cluster showed the most highly significant enrichment of pathways related to cell growth regulation (GO:0040009). Among the most highly upregulated epi_2 genes were the cell growth regulators *Malat1* (p = 3.35e-21) and *Notch2* (p=6e-3), the latter implicated in ChP hyperplasia and subsequent development of hydrocephalus in mice^130^. Co-immunostaining with antibodies directed against the ChP epithelial cell marker Otx2^130^ and the cell proliferation marker MKi67 confirmed the presence of ChP epithelial hyperplasia in *Pten* cKO animals vs. controls as early as P5 (**Fig. 4P, 4Q**).

ChP inflammation can lead to increased ChP-mediated CSF hypersecretion and hydrocephalus via upregulation of the cotransporter and other K^+^ and Cl^-^ channels and transporters on the ChP epithelial cell apical membrane^103^. The bumetanide sensitivity of the CSF hypersecretory response in the context of *Pten* cKO hydrocephalus implicates Nkcc1, the target of bumetanide^131,132^, in disease pathogenesis. Nkcc1 is a secondary active electroneutral Cl^-^ transporter that relies on gradients of Na^+^ and K^+^ ions for its activity^131,132^. Gene Ontology (GO) analyses of the most differentially expressed genes in the epi_2 sub-cluster also showed significant enrichment of pathways related to the transport of ions and carboxylic acid (**Fig. 4P, 4R**). Among the most highly upregulated epi_2 genes were *Slc12a2* (p = 3.41e-3), encoding Nkcc1, and *Kcnq1* (p = 5.30e-3), encoding the voltage-gated K^+^ channel Kv7.1.

Nkcc1 phosphorylation at its amino-terminal cytoplasmic domain threonines (Thr) Thr203, Thr207, and Thr212 by the serine-threonine kinase Spak is required for Nkcc1 activity^133^, and phosphorylation of the Spak-Nkcc1 complex at the apical ChP membrane can serve as indicator of ChP CSF secretory capacity^103,113^. TNFα and IL-1b stimulate Spak kinase activity in a Tlr4- and Nf-κB-dependent manner to increase ChP-mediated CSF production and epithelial transport in other epithelia^134,135^. Immunohistochemistry with antibodies directed against the phosphorylated, activated species of Spak (pSpak) and Nkcc1 (pNkcc1)^133,136-138^ showed significant upregulation of pSpak and pNkcc1 at the ChP apical membrane in both *Pten* cKO (**Suppl. Fig. 6**) and *Pten*^R335X/+^ mice (**Suppl. Fig. 6**) as compared to controls at P14. Notably, the pNkcc1 signal correlated with the extent of ventriculomegaly (**Suppl. Fig. 6**). These results suggest that an inflammation-induced increase in Nkcc1-mediated CSF secretion from a hyperplastic ChP contributes to the pathogenesis of *PTEN*-mutant ventriculomegaly.

### *Pten*-mutant ventriculomegaly is accompanied by ASD-like brain network dysfunction due to impaired activity of *Nkx2.1^+^* NSC-derived inhibitory interneurons

CH patients can exhibit persistence of ventriculomegaly and neurodevelopmental deficits despite CSF shunting, suggesting underrecognized impairment of intrinsic parenchymal brain function^13,14,139-141^. We hypothesized that *PTEN* mutation leads not only to hydrocephalus from aqueductal stenosis due to hyperproliferation of periventricular Nkx2.1^+^ neuroprogenitors, but also causes parenchymal circuit and network deficits due to altered activity of cortical inhibitory interneurons derived from these neuroprogenitors. As *Pten* deletion in MGE-derived cortical inhibitory interneurons negatively regulates the number of parvalbumin (PV+) interneurons in the somatosensory cortex and decreases the cortical interneuron population^142,143^, we examined cortical development and function of the somatosensory cortex in hydrocephalic P14 *Pten* cKO mice (*Pten*^flx/flx^; *Ai14 ^fl/wt^*; *Nkx2.1-*Cre). We confirmed reduced numbers of cortical tdTomato-positive cells at P14 in *Pten* cKO mice (**Fig. 5B**). However, quantitation of parvalbumin-expressing (PV+) neurons demonstrated a relative increase in the number of PV^+^ cells, yielding a greater proportion of tdTomato-positive interneurons (**Fig. 5C**).

**Figure 5.**
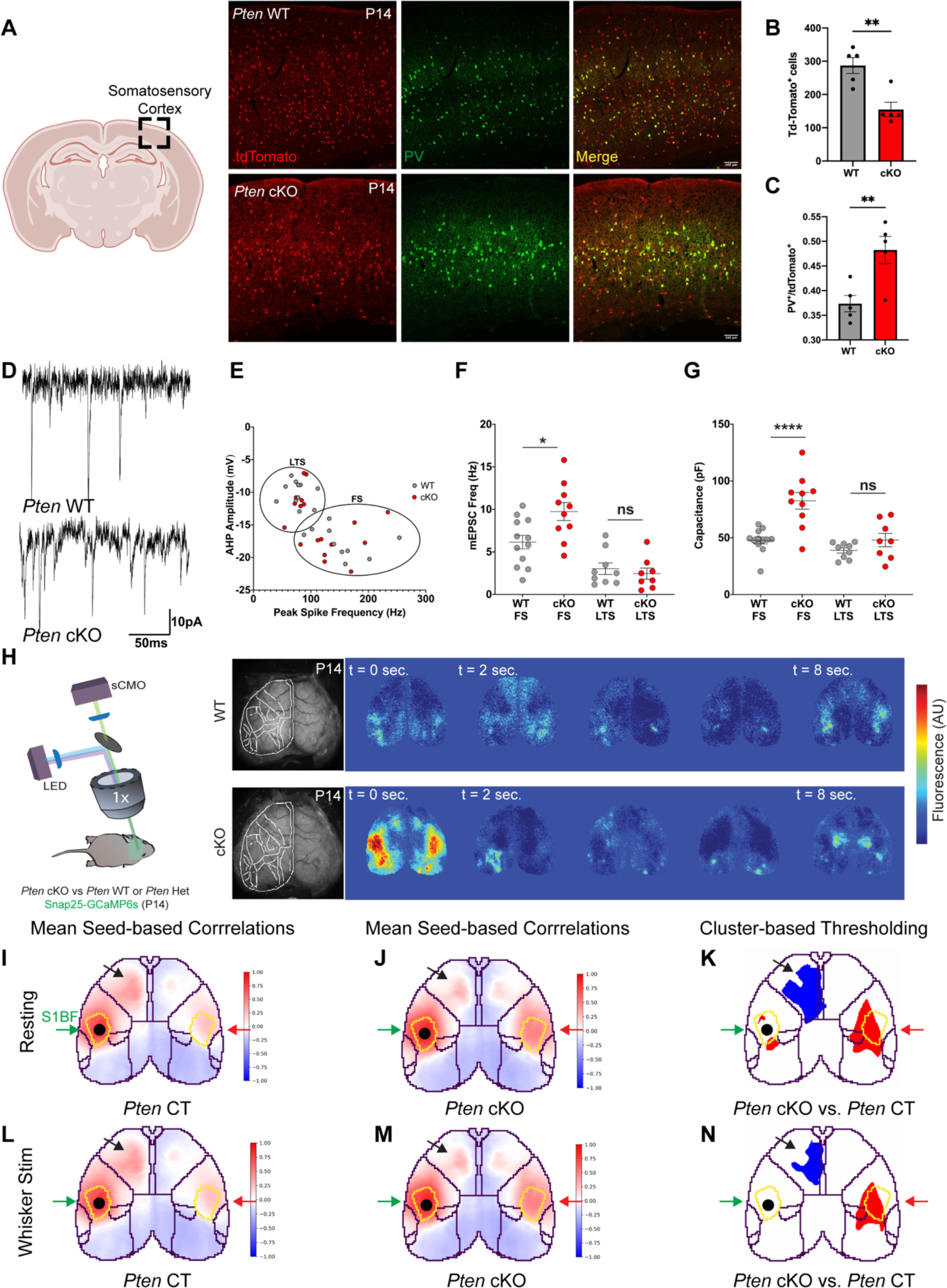
*PTEN*-mutant ventriculomegaly is accompanied by ASD-like network deficits in the somatosensory cortex, with activated Parvalbumin-positive cortical interneurons derived from the Nkx2.1^+^ neuroprogenitor lineage. (A) Left, schematic of mouse brain coronal section indicating somatosensory cortex. Representative somatosensory cortex immunofluorescent labeling of *Nkx2.1-Cre* lineage interneurons (tdTomato; red) and Parvalbumin (PV; green), in *Pten* cKO and *Pten* WT littermates, with quantitation of the number of tdTomato-positive cells (B) and of the ratio of Nkx2.1-lineage interneurons expressing Parvalbumin (C). Scale bars, 50 µm. Two-sided, unpaired *t*-test. (D) Representative traces of mEPCs recorded from tdTomato positive interneurons in the somatosensory cortex of layer II/III from *Pten* WT (above) and *Pten* cKO littermates (below).(E) *Nkx2.1-Cre*-lineage, tdTomato-positive interneurons from WT and cKO littermates, clustered by peak spike frequency as a function of afterhyperpolarization (AHP) amplitude into Fast Spiking cells (FS, putative parvalbumin-expressing cells) and low threshold spiking cells (LTS, putative somatostatin-expressing cells). (F) mPESC frequency is increased in FS but not LTS cells from *Pten* cKO as compared to cells from Pten WT mice. Two-sided, unpaired *t*-test. (G) Increased capacitance of FS but not LTS cells from *Pten* cKO FS as compared to that from corresponding *Pten* WT cells. Two-sided, unpaired *t*-test. (H) Left, schematic of widefield mesoscopic [Ca^2+^] imaging of awake, resting P13-P15 *Pten* ckO mice and littermate controls (*Pten* WT or *Pten* HET) expressing SNAP25-GCaMP6s. Right, representative fields of view with associated pseudocolor images from fluorescence emission video recordings. (I)-(J) Mean seed-based Pearson’s correlation maps for awake, resting *Pten* CT mice and *Pten* cKO mice (J) with a seed in the left barrel field somatosensory cortex. Green arrows highlight the seed in the left S1 barrel field (S1BF), red arrows highlight contralateral S1BF, and black arrows highlight ipsilateral motor cortex. (K) Cluster-based inference multiple comparisons corrections of Welch’s t-test contrasted *Pten* cKO and *Pten* CT mean seed-based correlations maps. (L)-(M) Mean seed-based Pearson’s correlation maps for awake, whisker-stimulated Pten CT mice and Pten cKO mice (See Methods) with a seed in the left barrel field somatosensory cortex. Green arrows highlight the seed in the left S1 barrel field (S1BF), red arrows highlight contralateral somatosensory cortex, and black arrows highlight ipsilateral motor cortex. (N) Cluster-based inference multiple comparisons corrections of Welch’s t-test contrasted *Pten* cKO and *Pten* CT mean seed-based correlation maps. Error bars, mean ± sem; each symbol represents one animal. *p<0.05, **p<0.01, ***p<0.001, ****p<0.0001, ns = not significant.

To explore the functional impact of these findings, we compared whole-cell currents recorded from tdTomato-positive interneurons of the somatosensory cortex of *Pten* WT and *Pten* cKO mice (**Fig. 5D**). We first recorded passive membrane properties under voltage-clamp and then switched to current-clamp to measure firing frequency and amplitude of after-hyperpolarization (AHP) amplitude. We used peak spike frequency and AHP amplitude to group the tdTomato^+^; *Nkx2.1*-Cre neurons as fast-spiking (FS) putative PV+ interneurons versus low threshold-spiking (LTS) putative somatostatin-positive neurons (**Fig. 5E**). We then examined miniature excitatory post-synaptic currents (mEPSCs) onto these interneurons to determine if *Pten* knockout increased excitatory drive. mEPSC frequency and capacitance were increased in *Pten* cKO FS cells (**Fig. 5F**, 4G, p<0.05, p<.0001); in contrast, mEPSC in LTS cells did not change significantly (**Fig. 5F, 5G**). These data suggest an increased proportion of PV+ interneurons receiving increased excitatory drive, consistent with previous results showing increased inhibitory drive on layer 2/3 pyramidal neurons in the somatosensory cortex of *Pten^flx/flx^; Nkx2.1*-cre+ mice, backcrossed into the CD-1 background^142^.

Early patterned neural activity is essential for the proper development of cortical connectivity^144^. In typically developing rodents, during the second postnatal week, pyramidal neurons in the primary somatosensory cortex (S1) undergo an increase in connectivity to ipsilateral primary motor cortex (M1) and a decrease in connectivity to contralateral S1^145-147^. Alterations of early activity disrupts this refinement^147,148^. Given that interneurons are critical regulators of patterned brain activity,^149^ we hypothesized that cortical connectivity would be altered in *Pten* cKO mice, which has never been examined in the context of *PTEN* mutation or hydrocephalus.

We therefore performed wide-field mesoscopic Ca^2+^ imaging of the cortical mantle of resting, awake, hydrocephalic *Pten* cKO mice and their littermate controls genetically-engineered to express SNAP25-GCaMP6s at P14 (**Fig. 5H**, see Methods). Using activity in the left somatosensory barrel field as a seed, we compared seed-based correlation maps for *Pten* cKO and control littermates. After correcting for multiple comparisons using cluster-based inference^150,151^, three clusters of altered connectivity emerged: (i) a positive cluster overlapping with the seed region indicating intrahemispheric hyperconnectivity within the seed region; (ii) a positive cluster overlapping with the contralateral seed region indicating interhemispheric hyperconnectivity, and (iii) a negative cluster in the ipsilateral motor cortex indicating intrahemispheric hypoconnectivity (**Fig. 4I-4K**).

Next, we examined functional connectivity in whisker-stimulated *Pten* cKO mice and littermate controls (see Methods), and found hyperconnectivity within the contralateral somatosensory seed region, and hypoconnectivity in the ipsilateral motor cortices, mirroring the results of the resting state analysis **(Fig. 4L-4N**). Similar circuit and network-level deficits have been reported in mouse models of Fragile X syndrome^152,153^. These results show *Pten*-mutant hydrocephalus is accompanied by ASD-like parenchymal network dysfunction due to impaired activity of *Nkx2.1^+^* NSC-derived inhibitory interneurons.

### Early post-natal everolimus treats *Pten*-mutant hydrocephalus and parenchymal brain deficits by rescuing mTORC1-dependent Nkx2.1^+^ cell pathology

PTEN negatively regulates mTORC1 activity^34^. Raptor is a critical component of mTORC1 complex activation and phosphorylation of 4EBP1^154,155^. We hypothesized Raptor deletion would inhibit mTORC1 activation caused by *Pten* knockout, thus attenuating development of ventriculomegaly in *Pten* cKO mice. To test this, we crossed in a *Raptor^flx/flx^* allele into *Pten*^flx/wt^; *Nkx2.1*-Cre mice to conditionally delete *Raptor* in *Nkx2.1-Cre*-expressing cells with concomitant *Pten* depletion. Strikingly, *Pten*^flx/flx^; *Raptor*^flx/flx^; *Nkx2.1-Cre* double knockout (“*Pten* <u>d</u>KO”) mice exhibited complete rescue of hydrocephalus compared to their *Pten*^flx/flx^; *Raptor*^flx/wt^; *Nkx2.1*-Cre control littermates (*Pten* cKO; *Raptor* cHet) at P14 (**Fig. 6D**, p<.01). Compared to controls, *Pten* dKO mice also had reduced caliber of the d3V area (**Fig. 6F**, p<.001) and reduced p-NKCC1 immunofluorescence at the ChP (**Fig. 6H**, p<.05). Finally, examination of survival out to P50 revealed *Pten*^flx/flx^; *Raptor*^flx/flx^; *Nkx2.1-Cre (Pten* dKO mice) prolonged survival of *Pten* dKO mice as compared to *Pten^f^*^lx/flx^; *Raptor*^flx/wt^; *Nkx2.1-Cre* mice. (**Fig.6B**, p<.0001). These data suggest that ventriculomegaly and altered CSF dynamics caused by loss of *Pten* is dependent upon Raptor-mediated activation of mTORC1.

**Figure 6.**
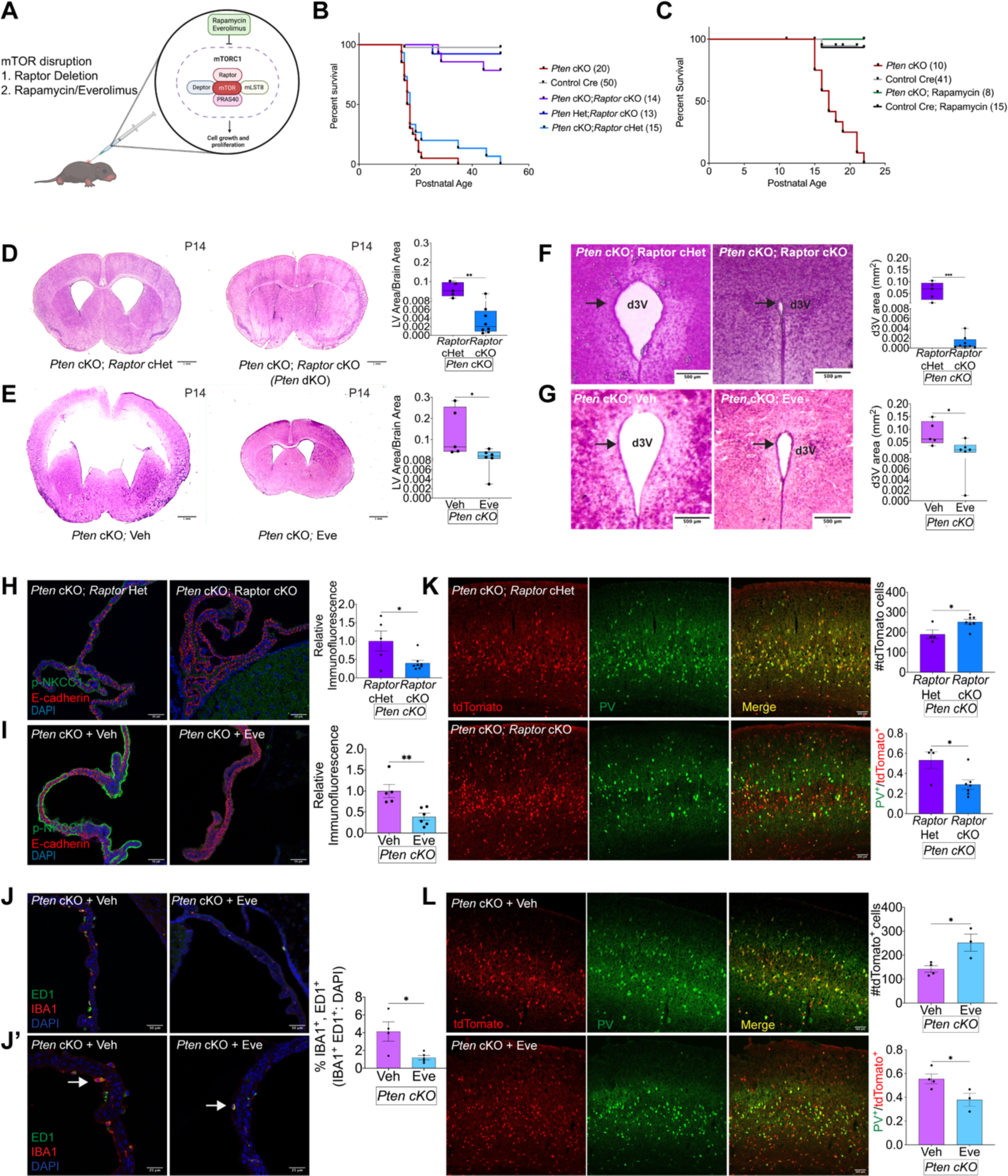
Genetic or pharmacological mTOR inhibition corrects ventriculomegaly and rescues lethality in hydrocephalic *Pten* mice. (A) Schematic illustrating strategy of mTOR inhibition with genetic deletion of *Raptor* or treatment with Everolimus (1 mg/kg body weight) every 48 hours from P1 to P14.(B) Kaplan-Meier survival curves of *Pten* cKO mice, *Pten* cKO; *Raptor* cKO mice, and their littermate control mice: Control Cre (Pten WT or Pten Het; Nkx2.1-Cre), *Pten* cKO; *Rator* Het, and *Pten* Het; *Raptor* Het mice. (C) Kaplan-Meier survival curves of *Pten* cKO mice, Control Cre (*Pten* WT or *Pten* Het; *Nkx2.1-Cre*), Control Cre mice treated with rapamycin (10 mg/kg body weight) from P10-P25, and *Pten* cKO mice treated with rapamycin (10 mg/kg body weight) from P10-P25. (D) Representative hematoxylin & eosin-stained coronal brain sections from P14 Pten^flx/flx^; Raptor^flx/wt^; *Nkx2.1-Cre* (*Pten* cKO; *Raptor* Het) and littermate Pten^flx/flx^; Raptor^flx/flx^; *Nkx2.1-Cre* (*Pten* cKO; *Raptor* cKO) mice with quantitation of total brain area-normalized lateral ventricular area. Scale bars, 1000 µm. Significance was tested by two-sided, unpaired *t*-test. (E) Representative hematoxylin & eosin-stained coronal brain sections from P14 Pten cKO mice treated with vehicle or everolimus, with quantitation of total brain area-normalized lateral ventricular area. Scale bars, 1000 µm. Two-sided, unpaired *t*-test. (F) Representative hematoxylin & eosin-stained coronal sections through dorsal third ventricle (d3V) from P14 *Pten* cKO; *Raptor* Het and littermate *Pten* cKO; *Raptor* cKO, with quantitation of ventricular normalized to total brain area. Scale bars, 500 µm. Two-sided, unpaired *t*-test.(G) Representative hematoxylin & eosin-stained coronal sections through dorsal third ventricle (d3V) from P14 *Pten* cKO mice treated with vehicle or with everolimus from P1-P14, with quantitation of ventricular area. Scale bars, 500 µm. Two-sided, unpaired *t*-test. (H) Representative ChP immunofluorescent labeling of pNKCC1 (green) and E-cadherin (red) in P14 *Pten* cKO; *Raptor* Het, and *Pten* cKO; *Raptor* cKO mice, with quantitation of p-NKCC1 intensity. Scale bars, 50 µm. Two-sided, unpaired *t*-test. (I) Representative ChP immunofluorescent labeling of pNKCC1 (green) and E-cadherin (red) in P14 *Pten* cKO treated with vehicle or with everolimus from P1-P14, with quantitation of p-NKCC1 intensity. Scale bars, 50 µm. Two-sided, unpaired *t*-test. (J and J’) Two representative ChP immunofluorescent labelings of ED1 (green) and IBA1 (red) in P14 *Pten* cKO treated with vehicle or with everolimus from P1-P14, with quantitation of % of IBA1+/ED1+ cells. Scale bars, 50 µm. (K) Representative somatosensory cortex immunofluorescent labeling of Nkx2.1-lineage interneurons (tdTomato; red), Parvalbumin (PV; green), in *Pten* cKO; *Raptor* Het and in *Pten* cKO; *Raptor* cKO mice, with quantitation of number of tdTomato-positive cells and % of tdTomato-positive cells expressing parvalbumin. Scale bars, 50 µm. Two-sided, unpaired *t*-test. (L) Representative somatosensory cortex immunofluorescent labeling of Nkx2.1-lineage interneurons (tdTomato; red), parvalbumin (PV; green), in *Pten* cKO mice treated with vehicle or with everolimus from P1-P14, with quantitation of number of tdTomato-positive cells and % of tdTomato-positive cells expressing parvalbumin. Scale bars, 50 µm. Two-sided, unpaired *t*-test.

Non-surgical treatments for CH do not exist^10^. Analogs of the naturally occurring mTORC1 inhibitor, rapamycin, are in current use to treat patients with tuberous sclerosis complex or PTEN hamartoma tumor syndrome^156-162^. We treated *Pten* cKO and littermate controls with rapamycin (10 mg/kg of body weight) every 24 hours, starting at P10, and observed a prolonged survival of mice out to P25 (**Fig. 5C**, p<.0001). Inspired by these results, we switched to the more clinically-relevant rapamycin analog, everolimus, to examine its effects on the development of hydrocephalus. We subcutaneously delivered vehicle or everolimus (1 mg/kg of body weight) every 48 hours from P1-P14. *Pten* cKO mice treated with everolimus (*Pten* cKO + Eve) displayed a significant reduction in area of the lateral third ventricule, normalized to total brain area (**Fig.6E,** p<.05), a decrease in the caliber of the d3V (**Fig.6G**, p<.05), and a reduction in the p-NKCC1 ChP immunofluorescence (**Fig.6I,** p<.01) compared to *Pten* cKO treated with vehicle (*Pten* cKO + Veh). Everolimus is used for immunosuppression after solid organ transplantation^163^. We found activated immune cells, indicated by immunohistochemical ED1 and IBA1 co-positivity, were reduced in everolimus-treated *Pten* cKO mice vs. vehicle-treated *Pten* cKO mice (**Fig 6L,** p<.05). These results suggest post-natal everolimus may be a non-surgical treatment of *Pten*-mutant hydrocephalus by antagonizing an inflammation-induced increase in ChP-mediated CSF secretion.

Everolimus was recently shown to be safe and potentially efficacious for neurocognitive and behavioral deficits in a double-blind controlled clinical trial in patients with *PTEN*-associated ASD and neurodevelopmental disorders^164^. Rapamycin prevents the increase in soma size, migration, spine density, and dendritic overgrowth in *Pten* knockout dentate gyrus granule neurons^45,165^. We hypothesized that early postnatal everolimus treatment or *Raptor* deletion, in addition to treating *Pten*-mutant hydrocephalus, might also restore the *Pten*-mutant associated reduction of *Nkx2.1^+^*-lineage interneuron numbers in the brain parenchyma, thus preventing the increased proportion of parvalbumin-expressing neurons that contribute to ASD-like network defects (see **Fig. 5A**). Contemporaneous deletion of *Raptor* and *Pten* prevented the reduction of *Nkx2.1^+^*-lineage cortical interneuron numbers (**Fig. 6M**, p<.05) and reduced the percentage of those expressing parvalbumin (**Fig. 6M,** p<.05) in P14 *Pten* dKO mice as compared to controls. Similarly, everolimus-treated *Pten* cKO from P1-P14 prevented the reduction in interneuron number (**Fig.5N**, p<.05) and the increase in the percentage of PV^+^ interneurons (**Fig. 6N**, p<.05) as compared to vehicle-treated *Pten* cKO mice. These results suggest that *PTEN*-mutant hydrocephalus and accompanying intrinsic parenchymal brain deficits can be treated with a repurposed FDA drug (everolimus) by rescuing mTORC1-dependent Nkx2.1^+^ cell pathology.

## DISCUSSION

Many CH patients have ASD and other functional brain deficits not responsive to neurosurgical CSF shunting^12,139,140^, confounding discussions with patient families about prognosis and expected outcomes from surgery. Patients with ASD and other neuropsychiatric conditions – if they have undergone brain imaging – can have perplexing enlargement of their cerebral ventricles, prompting neurosurgical evaluation, burdensome diagnostic procedures, and even unnecessary shunt placement. *PTEN* is one the most frequently mutated genes in ASD; here, we have shown that *de novo* mutations in *PTEN* are also highly associated with the development of ventriculomegaly and neurosurgically-treated CH. We found *PTEN* is highly expressed in human fetal *Nkx2.1^+^*NPCs of the subventricular MGE and their cortical interneuron descendants. ASD- and CH-associated *Pten* mutation or conditional *Pten* deletion in murine fetal *Nkx2.1^+^*NSCs phenocopies human ventriculomegaly, which results from aqueductal stenosis secondary to periventricular *Nkx2.1^+^* NSC hyperproliferation and CSF hypersecretion from inflammation-induced ChP hyperplasia. Ventriculomegaly is accompanied by ASD-like brain network dysfunction arising from impaired activity of *Nkx2.1^+^* NPC-derived inhibitory interneurons. Embryonic *Raptor* deletion or post-natal everolimus treats *Pten*-mutant ventriculomegaly, improves parenchymal brain deficits, and increases survival by antagonizing mTORC1-dependent *Nkx2.1^+^*cell pathology. These data suggest developmental pleiotropy of PTEN signaling in CH and ASD pathogenesis may originate in NPCs at the prenatal brain-CSF interface^166^.

Neurodevelopmental disorders are thought to originate from environmental and genetic insults during critical periods of fetal brain development. We therefore used transcriptomic data from the fetal human brain to assess potential spatial and temporal vulnerabilities of *PTEN* expression. We show that *PTEN* and *PI3K* signaling genes are most highly expressed during peak midfetal neurogenesis (PCWs 13-16) in neuroprogenitor cells (NPCs), newborn neurons, and cortical interneurons derived from the Nkx2.1^+^ NPC lineages within the SVZ of the MGE. The recapitulation of ventriculomegaly seen in mice with conditional *Pten* deletion limited to NPCs expressing *Nkx2.1* and their lineage (*Pten* cKO mice) highlights Nkx2.1^+^ NPCs as a critical locus for PTEN-related neurodevelopmental disorders and supports an NPC paradigm of CH^30^. Indeed, abnormalities in NPC proliferation have been previously reported in CH mouse models^167,168^. Knockout of Dusp16, a dual-specificity phosphatase that negatively regulates the MAPK growth pathway, also causes embryonic NPC niche expansion and subsequent development of obstructive CH^167^. We show that *Pten* cKO mice with severe hydrocephalus also demonstrate CSF obstruction due to brain overgrowth caused by over-expansion of the MGE with increased proliferation of *Nkx2.1^+^* NPCs. The coincident increase of NPC within the embryonic MGE co-labeled with mitotic markers and mTOR pathway activation in *Pten* cKO mice indicate mTOR-activated proliferation, suggesting brain overgrowth caused by dysregulated NPC proliferation as a common mechanism in obstructive CH.

In the absence of CSF obstruction, ventricular enlargement can also be caused by impaired CSF circulation by ependymal cilia or the overproduction of CSF from a hyperplastic^94,95,106^ or inflamed ChP^113^. Indeed, we showed ventriculomegalic *PTEN* patients and *Pten* mutant mice without CSF obstruction. In hydrocephalic *Pten* cKO mice, we found functional and morphologically matured ependymal cilia, which diminishes the likelihood of impaired motile cilia function as a contributing factor to development of ventriculomegaly. Moreover, *Pten* cKO mice exhibit ∼3 fold increase CSF production accompanied by immune cell activation and ChP hyperplasia, despite lack of *Nkx2.1-Cre* expression and preserved PTEN expression in the ChP of these particular *Pten* cKO mice. Interestingly, neuron-specific *Pten* deletion in mice has been shown to cause microglia/macrophage activation in the cortex, cerebellum, and hippocampus.^169^ Inflammation has become increasingly recognized as an exacerbating factor at the intersection of the gene-environment predisposition towards neurodevelopmental disorders^170^. Thus, neuroinflammation caused by *in utero* injection of low-dose lipopolysaccharide (LPS) exaggerated brain overgrowth and ASD-associated behaviors in a mouse model of *Pten* heterozygosity.^171^ Indeed, recent studies have reported inflammation-induced enlargement of the choroid plexus in schizophrenia^172^ and multiple sclerosis^173^; however, inflammation and growth of the choroid plexus in the context of CH or ASD has not yet been extensively examined.

We replicated and extended recent findings of enlarged choroid plexus volume and ventricular volumes in individuals with ASD as compared to neurotypical individuals using large-scale brain imaging databases^87,88^. We also performed the first scRNAseq analysis of the choroid plexus in a mouse model of congenital hydrocephalus. This analysis revealed up-regulated genes in the choroid plexus of *Pten* cKO mice related to activation of non-macrophage immune cells (*S100a8*, *S100a9*, *Lcn2*, *Stfa2*, and *C3*), and activation of macrophages (*Ccl2*, *Spp1*, and *Gpx3*). Of note, increased expression of the proinflammatory cytokine *Ccl2* was previously reported in CSF of individuals with ASD^84^, and of mice following maternal immune cell activation at the ChP^116^. Further, we show significant activation of the Spp1-IGF ligand-receptor pathway between the immune and epithelial cell clusters, which may underlie inflammation-dependent growth of choroid plexus epithelial cells. Indeed, we report up-regulated genes in ChP epithelial cells related to cell growth (*Malat1, Notch 2*) and ion transport (*Slc12a2*, *Kcnq1*). We also detected robust phospho-activation of *Slc12a2*-encoded NKCC1 and of SPAK, both previously associated with CSF-hypersecretion in animal models of inflammation-induced hydrocephalus^103^. The results together suggest that ChP inflammation may cause both hyperplasia and increased SPAK-NKCC1-mediated secretory capacity of the ChP.

Associations between ChP NKCC1 activity and brain inflammation have been recognized with increasing frequency^174^. A recent study in mice showed that systemic, administration of the NKCC1 inhibitor bumetanide attenuated brain inflammation caused by intracortical injection of LPS^174^. NKCC1 has also been implicated in many neurodevelopmental disorders, including ASD, and bumetanide has previously been shown to attenuate ASD severity in multiple clinical trials for ASDs^175-177^. However, a recent large-scale phase 3 clinical trial of bumetanide treatment in individuals with ASDs failed to show clinical efficacy. A possible explanation, considering ASD heterogeneity, may be that bumetanide treatment benefits only a subtype of ASDs. In support of this idea, elevated blood cytokine levels were recently found to predict responsiveness to bumentanide in patients with ASD. Thus, our findings suggest that increased ChP volume and ventriculomegaly detectable by non-invasive brain imaging may serve as structural biomarkers stratifying ASD-associated CH for bumetanide responsiveness. Of note, bumetanide has been previously shown to improve ASD symptoms in individuals with Tuberous Sclerosis Complex^178^.

Investigations of PTEN loss-of-function mouse models of ASD have focused primarily on development and function of excitatory neurons. However, our findings showing highest *PTEN* expression within inhibitory neuroprogenitors, newborn neurons, and cortical interneurons derived from the medial ganglionic eminence (MGE) indicate that PTEN has an important role in the development and function of inhibitory neurons, highlighting an underappreciated function of PTEN in development of neural circuits. Indeed, we confirm previous findings that Pten depletion in Nkx2.1+, MGE-derived neuroprogenitors causes an increase in the proportion of parvalbumin-positive cortical interneurons. We extend these findings with the discovery that disruption of mTOR pathway activation restored the number and proportion of parvalbumin-positive cells to wild-type levels. We further find that PTEN depletion selectively increased excitatory inputs onto parvalbumin-positive interneurons. Strikingly, Pten cKO mice have hypersynchronized network activity within the somatosensory cortex and desynchronized network activity between the somatosensory and motor cortices. The evidence of hypersynchronous activity is, moreover, consistent with the idea of a parvalbumin-activated network^179,180^. Indeed, optogenetic activation of parvalbumin positive interneurons within the S1 barrel cortex was recently shown to hypersynchronize network activity due to synchronized post-inhibitory rebound firing of pyramidal neurons^181^. Sensory dysfunction, a DSM-V criteria for ASD^182^, and motor delay has been reported in individuals with ASD harboring *PTEN* mutations^183^; likewise, sensory processing and sensorimotor gating deficits have been reported in Pten loss-of-function mouse models of ASD^42,184^. These findings together suggest that altered parvalbumin-positive neuron activity may contribute to cognitive and behavioral deficits of *PTEN*-mutated ASD by influencing fidelity of cortical somatosensory information flow.

Our study has demonstrated a therapeutically targetable association of ventriculomegaly with altered brain function. We showed that mTORC1 inhibition via conditional Raptor deletion in Nkx2.1+ NPCs completely rescues ventriculomegaly and parenchymal brain interneuronopathy caused by Pten depletion. To our knowledge, this is the first demonstrated genetic rescue of congenital hydrocephalus. This data is consistent with previous findings showing that mTOR deletion leads to reduced MGE size^79^, and that mTOR activation causes selectively increased connectivity and excitatory input to parvalbumin+ interneurons^185^. We also showed that early post-natal pharmacological inhibition of mTOR increased survival, reduced ventriculomegaly, and prevented interneuron pathology in *Pten* cKO mice. While the ventricles remained slightly enlarged in everolimus-treated Pten cKO mice, significant reduction in p-NKCC1 and immune cell activation at the ChP suggests reduced CSF production as a mechanism of this effect.

The use of rapamycin analogs such as everolimus to attenuate ventriculomegaly and correct parenchymal brain deficits has potentially high translational value. First, everolimus may be used as an adjunct therapy to neurosurgical CSF diversion to prevent neuronal dysfunction inherent in CH patients harboring mutations within the PTEN/PI3K/MTOR signaling pathway. Whole exome sequencing may thus be useful to stratify patients with CH for treatments to optimize neurodevelopment, as well as for post-surgical prognostication of neurodevelopment. Second, the recent development of brain-restricted mTOR inhibition could be exploited to target brain-specific mTOR pathology without side effects of somatic growth restriction^186^. To that end, detection of ventriculomegaly or ChP hyperplasia by neonatal brain imaging could serve as a radiographic “biomarker” to initiate preemptive brain-restricted mTOR inhibition to reduce or prevent ASD symptomatology without risking growth restriction. Third, our work highlights the need to start early behavioral intervention programs in CH upon initial diagnosis, and to implement early routine structural and functional brain MRI in ASD to improve both diagnosis and, ultimately, prognosis.

### Limitations of the study

This study demonstrates mTORC1 as the principal driver of cortical pathology in developing brain structure and function in Pten-mutated CH. However, the contribution of mTOR complex 2 (mTORC2) activation, which may also be inhibited by chronic rapamycin and its analogs^187^, to altered CSF dynamics and brain physiology in Pten mutants was not examined. Future investigations will examine brain structure and function In Pten cKO mice after the deletion of Rictor, the rapamycin-insensitive regulator of mTORC2 activation^188^. Furthermore, our functional studies primarily employed conditional deletion of Pten and Raptor in cells expressing Nkx2.1-Cre and their lineage; however, non-cell-autonomous effects may emerge in the setting of gene deletion in a large population of cells in brain tissue. As such, we cannot exclude the possibility of a non-cell-autonomous contribution of impaired CSF dynamics to our analyses of brain parenchymal interneuron development and altered functional connectivity in *Pten* cKO mice. Indeed, in mice it has been reported that factors present within CSF may cause precocious development of parvalbumin+ interneurons, also leading to a decreased number of parvalbumin+ interneurons in the cortex^189^. However, conditional Pten deletion within the Nkx2.1-Cre lineage in the CD1 background leads to selective parvalbumin+ interneuron abnormalities without grossly apparent alterations in CSF dynamics^142^. Future investigations will examine how choroid plexus function may independently cause altered development of interneurons and their function in the assembly of cortical circuits.

## METHODS

Key resources table

| REAGENT or RESOURCE | SOURCE | IDENTIFIER |
| --- | --- | --- |
| **Antibodies** |  |  |
| Pten | CST | Cat# 9188, RRID: AB_2253290 |
| p-Nkcc1 | Millipore Sigma | Cat# ABS1004 |
| Iba1 | Abcam | Cat# ab5076, RRID:AB_2224402 |
| Ed1/CD68 | Cell Signaling | Cat# 79594, RRID:AB_2799935 |
| E-cadherin | BD Biosciences | Cat# 610181, RRID:AB_397580 |
| Nkx2.1 | Abcam | Cat# ab76013, RRID:AB_1310784 |
| Lhx6 | Santa Cruz | Cat# sc-271433, RRID:AB_10649856 |
| pH3-488 | CST | Cat# 3465, RRID:AB_10695860 |
| tdTomato | Origene | Cat# AB8181, RRID:AB_2722750 |
| p-4EBP1 | CST | Cat# 2855, RRID:AB_560835 |
| Parvalbumin | Synaptic Systems | Cat# 195 004, RRID:AB_2156476 |
| p-SPAK | Millipore Sigma | Cat# 07-2273, RRID:AB_11205577 |
| Otx2 | R&D Systems | Cat# AF1979, RRID:AB_2157172 |
| Ki67 | Abcam | Cat# ab15580, RRID:AB_443209 |
| **Chemicals, peptides, and recombinant proteins** |  |  |
| Rapamycin | Cayman Chemical | Cat#13346 |
| Everolimus | Cayman Chemical | Cat#11597 |
| Evans Blue | Sigma Aldrich | Cat#E2129 |
| Hematoxylin & Eosin Stain Kit | Vector | Cat#H- |
| **Deposited data** |  |  |
| Raw and analyzed single-cell RNA-seq | This paper | GEO:XXXXX |
| **Experimental models: Organisms/strains** |  |  |
| *Pten*<sup>flx/flx</sup> | Jackson Laboratory | RRID: IMSR_JAX:006440 |
| *Raptor*<sup>flx/flx</sup> | Jackson Laboratory | RRID: IMSR_JAX:013188 |
| *Snap25-2A-GCaMP6s-D* | Jackson Laboratory | RRID: IMSR_JAX: 025111 |
| Pten R335X mouse model | This paper |  |
| **Software and algorithms** |  |  |
| Fiji software | Freely available online | https://imagej.net/Fiji |
| GraphPad Prism 9 | GraphPad software | https://graphpad.com/scientific-software/prism/ |
| Seurat | https://github.com/satijalab/seurat |  |
| R | https://www.r-project.org/ |  |
| MATLAB 9.3 | MathWorks | https://mathworks.com/products/compiler/matlab-runtime.html |

### Resource Availability

### Lead Contact

### Materials Availability

Data and Code Availability

Single-cell RNA-seq data have been deposited at GEO and are publicly available as of the date of publication. Accession numbers are listed in the key resources table. Any microscopy data, confocal, or other, reported in this paper will be shared by the lead contact upon request.

Python and MATLAB code to analyze calcium imaging data are available from Lead contact without restriction. All previously published algorithms are listed in the key resources table.

Any additional information required to reanalyze the data reported in this paper is available from the lead contact upon request.

### Experimental Model and Subject Details

#### Whole-exome sequencing and variant calling

Exon capture was performed on genomic DNA samples derived from saliva or blood using Roche SeqCap EZ MedExome Target Enrichment kit or IDT xGen target capture kit, followed by 101 or 148 base-paired-end sequencing on the Illumina platforms. BWA-MEM^190^ was applied to align sequence reads to human reference genome GRCh37/hg19. GATK HaplotypeCaller^191^ and Freebayes^192^ were used to call single-nucleotide variants and small indels, and ANNOVAR^193^ was applied to annotate. Allele frequencies annotated were sourced from data in the Exome Aggregation Consortium (ExAC)^194^, gnomAD (v.2.1.1) and Bravo databases^195,196^. Deleteriousness of missense variants was predicted using MetaSVM and MPC algorithms (D-Mis, defined as MetaSVM-deleterious or MPC-score ≥ 2). Inferred LoF variants consisted of stop-gain, stop-loss, frameshift insertions/deletions, canonical splice site and start-loss. LoF and D-Mis mutations were considered ‘damaging’^197^.

DNVs were called using an established TrioDeNovo-based pipeline^198^ and further filtered to variants in exonic or splice-site regions, with read depth (DP) of 10 in the proband and both parents, with minimum proband alternative read depth of 5, with proband alternative allele ratio ≥28% if having <10 alternative reads or ≥20% if having ≥10 alternative reads, with alternative allele ratio in both parents ≤3.5%, and with a global MAF ≤ 4 × 10−4 in the Exome Aggregation Consortium database as previously described^29^.

After filtering using the aforementioned criteria for each type of mutation, *in silico* visualization was performed to remove false-positive calls. Variants in the top candidate genes were further confirmed by Sanger sequencing.

#### *De novo* enrichment analysis

Enrichment for *de novo* variation was performed using DenovolyzeR v0.2.0^199^. The expected number of DNVs in the case and control cohorts across each functional class was calculated by taking the sum of each functional class-specific probability multiplied by the number of probands in the study x 2 (diploid genomes). Then, the expected number of DNVs across functional classes was compared to the observed number in each study using a Poisson test.

To examine whether any individual gene contains more protein-altering DNVs than expected, the expected number of protein-altering DNVs was calculated from the corresponding probability, adjusting for cohort size. The Poisson test was then used to compare the observed DNMs for each gene versus expected. As separate tests were performed for protein-altering, protein-damaging and LoF DNVs, the relevant Bonferroni multiple-testing threshold is therefore equal to α = 8.6 × 10^-7^ = (0.05/[3 tests x19,347 genes]).

#### Congenital hydrocephalus gene list composition

Due to the documented contribution of *de novo* variants to CH disease burden^4,29,200,201^, experimentally-determined risk gene lists for congenital hydrocephalus were composed based solely upon DNV enrichment analysis. The potential CH risk gene list (p-CH) was inclusive of any gene with a pLI ≥ 0.90 in ExAC which had at least one ‘damaging’ (LoF variant or missense variant with MPC ≥ 2 and/or MetaSVM = ‘D’) *de novo* variant. The high-confidence CH gene list (hc-CH) was inclusive of any gene with a pLI ≥ 0.90 in ExAC with at least two ‘damaging’ *de novo* variants. The exome-wide-significant CH gene list (EWS-CH) is inclusive of any gene which reached the threshold for exome wide significance (α = 8.6 × 10^-7^) in any of the three queried classes of variants. An extensive literature search was also conducted to compose a non-experimentally-determined CH risk gene list inclusive of genes documented as significantly associated with CH pathogenicity in the literature^200,201^ and was used for comparison against the experimentally-determined lists.

#### Gene ontology analysis

Using the identified CH gene lists and gene modules significantly enriched with the CH gene list, enrichment analysis was performed for gene ontologies (biological processes, cellular components, and molecular functions) and biological pathways (WikiPathways) using the EnrichR R package Version 3.0^202^. The top 10 terms with the lowest adjusted p-values were reported in the order from highest to lowest combined Z-scores. Adjusted p-values < 0.05 were considered significant.

#### DisGeNet analysis

Overlap analysis between the p-CH gene list and disease risk gene list curated by DisGeNET, one of the largest available databases documenting gene association with disease terms, was conducted applying the DisGeNET2R R package for DisGeNET v7.0^203^. Iterative one-tailed Fisher’s exact tests were applied to assay genetic overlap between the p-CH gene list and risk gene lists for all disease terms documented in DisGeNET as described previously [￼Cite AC paper here]. The analysis was corrected for false-discovery using the Benjamini-Hochberg method. Adjusted p-values < 0.05 were considered significant.

#### Developmental Human Brain ScRNAseq Dataset Analysis

As described previously^67^, the preprocessing and clustering analysis for scRNA developmental human brain dataset was completed using Seurat^204^. Briefly, cells with fewer than 1000 genes/cell were removed, as were cells with greater than 10% of their individual transcriptome represented in either mitochondrial or ribosomal transcripts. Only genes expressed in at least 30 cells were carried forward in the analysis. The raw counts were normalized and log2-transformed by first calculating ‘size factors’ that represented the extent to which counts should be scaled in each library. Highly variable genes were detected using the proposed workflow and subsequently used for unsupervised dimensionality reduction techniques and principal component analysis. UMAP coordinates were calculated using standard Seurat workflow, and clusters were assigned to cells based on previous analysis by a hybrid method using Louvain clustering and WGCNA^67^. The non-parameteric Wilcoxon rank sum test was used to identify differentially expressed markers across cell clusters and laminar zones by running FindAllMarkers. Heatmap expression values were calculated using AverageExpression function and visualization of the heatmaps were created using the pheatmap package

#### Spatio-Temporal cell Atlas of the human Brain (STAB) ScRNAseq Dataset Analaysis

As described previously^63^, Seurat was used to preprocess, integrate and cluster 13 scRNA-seq human brain dataset datasets and remove batch effects of multiple datasets. Briefly, cells with fewer than 1000 genes and genes expressed in less than 30 cells were removed. Each gene’s expression level was normalized and defined as the ratio between its counts and the total counts of genes expressed in the cell, which was then multiplied by a scale factor (10 000 by default) and log-transformed. Given a dataset, the top 2000 genes with the highest expression variance across cells were determined for downstream analysis. Then, using the integration tool of Seurat, the anchors representing pairwise cell correspondences between single cells across datasets were identified. These anchors were then used to transform distinct datasets into a shared space and construct an integrated scRNA-seq dataset. Violin plots were used to visualize gene expression patterns across time and brain region.

### Mouse experiments

All experiments were done in accordance with the regulations set forth by the Yale University animal care and use committee and per the guidelines and regulations in the NIH Guide for the Care and Use of Laboratory Animals.

#### *Pten* cKO and *Raptor* cKO mice

*Nkx2.1-cre* (Jackson Laboratory, Stock No: 008661) mice were mated with *Pten^fl/fl^* (Jackson Laboratory, Stock No: 006440), Cre-dependent tdTomato expression locus (Jackson Laboratory, Stock No: 007914), and Raptor^fl/fl^ (Jackson Laboratory; Stock No: 013188) to obtain *Nkx2.1-Pten^fl/fl^;Ai14* mice in which Pten and Raptor are conditionally deleted from embryonic NSCs in the ventral telencephalon, hypothalamus, pituitary, lungs, and thyroid gland. Male and female animals were randomly selected and used in all investigations of this study. All strains are congenic on the C57BL/6J genetic background. *Pten^R335X^* mutant mice were generated as described below.

#### CRISPR/cas9 generation of *Pten* R335X mutant mice

Pten point mutation R335X mice were generated by the Yale Genome Editing Center. Mutation of Pten R335 to a termination codon (CGA->TGA) was performed via CRISPR/Cas9-mediated genome editing essentially as described (Yang, Wang, and Jaenisch 2014; Chen et al. 2016). Potential Cas9 target guide (protospacer) sequences in the vicinity of the R335 CGA codon were screened using the online tool CRISPOR (http://crispor.tefor.net). Templates for sgRNA synthesis were generated by PCR, and sgRNAs were transcribed *in vitro* and purified using Megashortscript and Megaclear kits (Life Technologies). Cas9 enzyme was purchased from NEB. Candidate sgRNA/Cas9 RNPs were complexed and tested by zygote electroporation, incubation of embryos to blastocyst stage, and genotype scoring by indel creation at the target site. The sgRNA containing the protospacer sequence ATTTGGAGAGAAGTATCGGTTGG was highly efficient and was used for mouse creation. sgRNA/Cas9 RNP was co-electroporated with a 127-bp repair oligonucleotide into C57Bl/6J zygotes. The asymmetric repair oligo (“reverse strand”) incorporates the reverse complement of the codon substituting R^335^ to X. The repair oligo sequence 5’ AGTATAGAGCGTGCAGATAATGACAAGGAGTATCTTGTACTCACCCTAACAAAAAACGATCTTGACAAAGCAAACAAAGACAAGGCAAACTAGTACTTCTCTCCAAATTTTAAGGTCAGTTAAAACC was synthesized by IDT. Electroporated zygotes were transferred to pseudo-pregnant CD-1 foster females using standard techniques (Behringer et al., n.d.). Genotyping was performed by PCR amplification followed by Sanger sequencing of a 324-bp fragment. Primers for PCR were PtenF 5’ TCTTCATACCAGGACCAGAGGAA and PtenR 5’ AAC AGA AAA GGA AAG GCT ACG CAA. Correct alleles were confirmed by TA cloning of PCR products and sequencing. 2 mice carrying the proper mutation were identified and crossed to C57BL/6J mice to demonstrate germline transmission. The line was then further backcrossed fiver generations to breed out any off-target mutations. All Pten R335X mice were genotyped by sequencing using PtenF and PtenR primers.

#### SNAP25-GCaMP6s mice

GCaMP6s (Jackson Laboratory, Stock No: 025111) mice, provided by Hongkui Zeng^205^, were mated with Nkx2.1-Cre; Pten^fl/wt^ mice as described above for widespread brain expression of GCAMP6s slow variant calcium indicator.

#### Mouse magnetic resonance imaging and quantitation of ventriculomegaly

Structural MR imaging data were collected on an 11.7T preclinical Bruker magnet (Bruker, Billerica, MA) using an in-house built saddle coil for optimal whole brain sensitivity. Mice were anesthetized with isoflurane (3% induction, 1-2% maintenance, in a medical air and oxygen mixture 1:0.5). During imaging, body temperature was maintained with a circulating water bath. A RARE (Rapid Acquisition with Relaxation Enhancement) imaging sequence was used. Over a period of 31 minutes and 30 seconds, using a TR/TE of 4500/11.25ms, 20 repetitions, and rare-factor 8, a coronal 0.07×0.30×0.07mm^3^ image of the whole brain (FOV 1.40×8.40×12.0mm^3^) was obtained. This sequence was repeated, and the data averaged after motion correction using customized modules within Bioimage Suite (BIS) Web (www.bioimagesuite.org) developed in-house and available free online. In post-processing, again using BIS, signal intensity thresholding was delineated in the ventricles from surrounding brain tissue, ventricular volume was measured, and 3D reconstructions were performed.

#### Histological quantitation of ventricular area

Brains from control and *Pten* mutant mice at P14 were postfixed in 4% paraformaldehyde for 24 hours, dehydrated in 30% PBS-sucrose solution for 48 hours at 4°C, embedded in optical cutting temperature solution, flash-frozen at −20°C, and then kept at −80°C until further use. 20mm coronal brain sections were cut with a Cryostat (Leica, cat# CM1950), and stained with hematoxylin and eosin according to the manufacturer’s protocol (Vector cat #H-3502). Sections were imaged using Leica IC90E camera mounted to Leica M60 stereomicroscope. The areas of the lateral ventricle and brain were measured at 0.5 cm anterior to the bregma using Image J. The ratio of ventricular area to brain area was calculated to determine degree of ventriculomegaly. The data was then calculated as means ± standard deviations and evaluated using the Student’s t-test in Prism version 9.

#### Histological staining and Immunofluorescent labeling of brain sections

Mice were sacrificed following Yale IACUC guidelines. Briefly, brains were isolated, and tissue fixed in cold 4% PFA in 1X PBS and incubated at 4C overnight, transferred to 30% sucrose in 1X PBS overnight. Tissue was embedded in OCT-filled molds (Optical Cutting Temperature, Fisher, Cat#4583), rapidly frozen on dry ice and kept at −80C until sectioning. 20 µm sections were cut with a Cryostat (Leica, cat# CM1950) and serially collected into a 24 well plate in 1X PBS. Prior to histological staining and immunolabeling, tissue sections were mounted onto glass slides (Fisher, cat# 22-037-247) and air dried. Immunofluorescent labeling was done by permeabilizing tissue in in PBS/TritonX 0.4% (PBS-T), antigen retrieved in Tris-EDTA (1 mM, pH 8), washed and blocked in 10% Donkey serum/PBS-T for 1 hour at room temperature, followed by primary antibody incubation in 5% Donkey serum/PBS-T at 4C overnight. All antibodies used are listed in Key Resources. Unbound antibodies were washed in PBST 3x 15min, fluorescent-conjugated secondary antibodies were incubated for 2 hours at room temperature (1:500 in blocking solution, Table 1), then washed in PBST 3x 10 min. Autofluorescence was quenched using TrueView with antigen-retrieved tissue (Vectors Lab, cat# SP-8400), and nuclei were counterstained with DAPI/PBS-T (5ng/ml) for 10min, washed PBST 3x 5min, air dried and mounted on a coverslip with VectaShield Vibrance Antifade mounting media (Thermo, cat# 152250). Specimens were cured in the dark at room temperature for 24 hours prior to image acquisition by a Zeiss 880 Confocal Microscope (Yale Imaging Facility Science Hill).

#### Cell Counting and statistical analysis

To measure proliferation in the MGE, cell counting using Image J was performed on confocal z-stack images (5µm) taken with a 20X objective (Zeiss LSM800) of 20 µm coronally-cryosectioned, immunolabelled tissues at equivalent rostro-caudal positions. Similar to as a previously described^142^, tdTomato+ cells per area was determined by counting the number of cells from confocal images taken with a 10x objective (.7 zoom), which included all layers of the neocortex, and dividing the counted cells by the area of the neocortex from the image (cells/mm ^2^) The percentage of tdTomato+ cells that were co-labeled with parvalbumin was calculated by dividing the number of co-labeled cells by the total number of tdTomato+ cells. The percentage of proliferative (Ki67+) ChP epithelial cells (Otx2+) and activated immune cells (Ed1+, Iba1+) were counted and divided by the total number of counted DAPI+ cells to normalize for differences in ChP cells in each image. Unpaired t-tests were performed using Prism version 9, with p values < 0.05 considered significant.

#### Intraventricular injection of Evan’s Blue Dye (CSF tracer) and IVIS imaging

Mice were placed in a stereotaxic frame and anesthetized with 1.5% isoflurane in 100% oxygen and a small burr hole was made over the right lateral ventricle. Next, 5 mL Evans Blue Dye (1% in sterile aCSF, Harvard Apparatus, cat#59-7316) was slowly infused (0.5 mL/minute) into the right lateral ventricle (coordinates, x=1.0mm, y= .1mm, z= 2.00 mm, relative to bregma). Mice were immediately sacrificed and brains dissected 20 minutes after dye infusion to assess distribution. After 24 hours in 4% PFA, brains were imaged with IVIS imaging system (Caliper, CA). Fluorescence intensity (AF 555 for tdTomato, and AF647 for Evans Blue CSF tracer) in each brain was captured with a camera and measured using Living Image 3.0 (Caliper, CA).

#### Measurement of CSF flow rate

Flow velocimetry was determined using a two-step process of automated detection and tracking of particles followed by velocity averaging. Automatic detection of particles and their trajectories was achieved using Mosaic Particle Tracker plugin build in ImageJ^206,207^. Manual region of interest (ROI) segmentation was performed using the MATLAB tool CROIEditor. All particle tracking data outside of the ROI was removed before further analysis. Trajectories were then averaged across all frames of each video to yield mean flow vectors at each pixel position. For visualization, average flow velocity was calculated for 10-by-10 pixel non-overlapping windows, excluding outliers above the 97th percentile which were also removed from further analysis. The number of particles was calculated as the mean number of particles detected across all frames and was normalized using the sample’s ROI area. All image and data processing was performed using Matlab (The MathWorks, Natick, Massachusetts, USA, Version R2019a) and ImageJ (National Institutes of Health, USA, Version 1.5.3c).

#### Measurement of CSF production rates

Rates of CSF production were measured using our published method^113^ and as used previously^102^. Briefly, anesthetized mice were mounted in a stereotactic apparatus and a small burr hole was made over the right lateral ventricle (coordinates, x=1.0mm, y= .1mm, z= 2.00 mm, relative to bregma). Next, the mouse’s head was gently rotated on the ear-bars 90ᵒ, nose-down, and the suboccipital muscles were dissected to the cisterna magna to expose the atlanto-occipital ligament. The ligament was punctured with a cannula, a 30-gauge needle (BD, Cat#305106) connected to polyethylene tubing (Intramedic, PE-10) and advanced 2 mm through the foramen of Magendie to the 4th ventricle and fixed in place to skull with cyanoacrylate adhesive (Covetrus, cat#031477) and dental cement (Person Dental, Unifast Trad). Sterile, molecular grade mineral oil (100 µL; Sigma Aldrich, St. Louis, MO) was infused with a syringe pump (Pump elite 11, Harvard Apparatus) into the 4th ventricle at .5 mL/min over 1 minute to occlude the aqueduct of Sylvius, thereby creating a closed system of CSF circulation. With the mice in the same position, another cannula (30-gauge needle connected to PE-10 tubing) was advanced through the burr hole into the right lateral ventricle (2 mm beyond entry) and fixed in place with cyanoacrylate adhesive (Covetrus, cat#031477) and dental cement (Person Dental, Unifast Trad). CSF egress from the right ventricle through the cannula was allowed to flow out of the cannula for 10 minutes to equilibrate CSF flow before quantitation of CSF flow rate. After 10 minutes, the position of CSF within the PE-10 tube cannulated to the right lateral ventricle was marked at 5 min intervals for 30 minutes to quantify CSF production. The volume (V) of CSF that had formed at a given time point was calculated as: *V* (*mm*^3^)=π·*r*^2^·*d*V mm^3^=π·r^2^·d, where r is the radius of the PE tubing and d is the distance CSF traveled within the tubing. The rate of CSF formation (nL/min) could be calculated from the slope of the volume–time relationship.

#### scRNAseq sample preparation

Animals were anesthetized with Ketamine/Xylazine and transcardially perfused with ice cold PBS before brain removal. Choroid plexi from bilateral lateral ventricles were microdissected in ice cold HBSS with 30uM Actinomycin D (ActD, Sigma, cat#A1410) and digested at 37°C for 20 min with Collagenases-I and - IV (10 U/mL, 400 U/mL final, Worthington, Cat# LS004194, and LS004184) and DNase I (30 U/mL, Roche, cat# <u>04716728001</u>) and ActD (3uM), using a protocol adapted from Van Hove et al(Van Hove et al., 2019). Cells were subsequently washed in FACS staining buffer (0.5% BSA, 2mM EDTA in PBS without Calcium/Magnesium) and centrifuged at 350 rcf for 5 minutes. Cells resuspended in 0.4% BSA/PBS and 3uM ActD were sent for 10X 3prime library preparation and sequencing at Yale Center for Genome Analysis.

#### scRNAseq data analysis

Preprocessing and integration of *Pten* WT and *Pten* cKO rat ChP scRNAseq datasets

The preprocessing and clustering analysis for scRNA rat choroid plexus cell datasets was completed using Seurat^204^. Cells with < 200 genes, <500 UMIs (500) or >10,000 UMIs and mitochondrial gene percentage greater than 10% were removed. The normalization and initial feature selection for each of the two datasets (*Pten* WT and *Pten* cKO) were completed individually. The filtered matrices were normalized using ‘LogNormalize’ methods in Seurat, using a size factor of 10,000 molecules for each cell. The 2,000 most variable genes were identified by variance stabilizing transformation for each dataset through implementation of the FindVariableFeatures function in Seurat v4 (selection.method = “vst”). The experimental conditions were integrated using Seurat’s integration pipeline. For integration, 2,000 shared highly variable genes were identified using Seurat’s ‘SelectIntegrationFeatures()’ function. Integration anchors were identified based on these genes by canonical correlation analysis using the ‘FindIntegrationAnchors()’ function. The data were then integrated using ‘IntegrateData()’ and scaled again with ‘ScaleData()’. The integrated dataset consisted of 1896 *Pten* WT and 3302 *Pten* cKO cells.

Principal component analysis (PCA) and uniform manifold approximation and projection (UMAP) dimension reduction with 50 principal components were performed. A KNN graph was constructed based on the euclidean distance in PCA space to embed cells in a graph structure, and the edge weights were refined between any two cells based on the shared overlap in their local neighborhoods (Jaccard similarity). This step was performed using the FindNeighbors (reduction=’pca’, dims=1:50) command by taking the first 50 principal components as input. To cluster the cells in an unsupervised fashion, the Louvain algorithm (default) was applied to iteratively group cells together, with the goal of optimizing the standard modularity function. The FindClusters(resolution=0.3) function was implemented for this procedure. For visualization purposes, dataset dimensionality was further reduced to 2D embeddings using UMAP on the significant PCs via RunUMAP() functions of the Seurat package in R (Becht et al., 2018). The non-parameteric Wilcoxon rank sum test was used to identify differentially expressed markers across all clusters by running FindAllMarkers(dataset, only.pos = TRUE, min.pct = 0.25, logfc.threshold = 0.25). Further sub-clustering of immune cell populations was conducted in an unsupervised fashion using the Louvain algorithm. VlnPlot and DotPlot functions were used to visualize gene expression profiles across clusters and conditions.

To infer cell–cell interactions based on expression of known ligand–receptor pairs in different cell types, CellChat^119^ was applied to our experimental conditions using the official workflow and databases. Briefly, the normalized counts were loaded into CellChat, after which the preprocessing functions identifyOverExpressedGenes, identifyOverExpressedInteractions and projectData with standard parameters set were applied to our datasets. The main analyses were conducted using the functions computeCommunProb, computeCommunProbPathway and aggregateNet with fixed randomization seeds.

#### Cluster abundance analysis

To test the difference in cell counts between *Pten* WT and *Pten* cKO, we used beta-binomial generalized linear model in package aod::betabin. In our generalized linear model, we set the count of the cell type of interest and the total count of cells of each experimental condition to be the response variable and the experimental condition of the cells to be the independent variable. We tested for Pearson’s correlation between the frequency of each cell cluster. We adjusted the *p* value threshold using Bonferroni correction (0.05/20 = 0.0025).

#### Gene Ontology (GO) enrichment analysis

The gene lists for differentially expressed genes from cell clusters of *Pten* WT and *Pten* cKO ChP ScRNA-seq datasets were studied for gene ontology, pathway and upstream transcription factor enrichment using the Enrichr R package^208^. Enrichr contains a diverse and up-to-date collection of over 100 gene set libraries available for analysis and download. The databases for these analyses include gene ontologies (biological processes, cellular components, and molecular functions), biological pathways (Wiki pathways Human and Mouse). Adjusted p-value of less than 0.05 was considered significant and combined Z-score was used for ranking.

#### Lateral ventricle ChP quantitation

As previously described^209^, semi-quantitative analysis of protein abundance in the CPe was performed by measuring immunofluorescence intensities from confocal images taken with a 20X objective (Zeiss 880). Experimental and control tissues were processed in parallel, using uniform parameters during immunolabeling and imaging. Fixed settings (laser power, gain, image depth, offset, and averaging) were used while acquiring confocal images in the focal plane with the highest signal intensity for all images with a given antibody. Image J was then used to quantify immunofluorescence, where the region of interest was manually selected for measurement, which was then divided by the number of cell nuclei within the region of interest to normalize for differences in ChP cell number among images. The data were then normalized to the mean of the controls’ fluorescence signal to generate relative fluorescence intensities. Unpaired t-tests were performed using Prism version 9, with p values < 0.05 considered significant.

#### Human magnetic resonance imaging and quantitation of ventriculomegaly

Non-contrast T1 MRI imaging was acquired from the Autism Brain Imaging Data Exchange (ABIDE) collections^87,88^ for 1051 subjects with ASD and 1163 controls. By applying SynthSeg+, a robust convolutional neural network for brain imaging segmentation^89^, all subjects’ imaging was resampled in 1mm isotropic space and ventricles were delineated, facilitating the calculation of ventricular volumes and total intracranial volumes. A total of 920 subjects with ASD and 1003 controls passed SynthSeg+’s automated internal quality control metric (QC>0.65). Ventricular volume, represented as percent of total intracranial volume, was compared between ASD and controls by ANOVA controlling for subject sex and age at scan. FastSurferCNN was applied to segment these subjects’ resampled MRIs^90^ and choroid plexus volume was also compared between ASD and controls by ANOVA controlling for subject sex and age at scan. For visualization, subject MRIs were affine transformed to MNI152 2009b NLIN asymmetric space^210^ using Advanced Normalization Tools (ANTs)^211^ and ventricular three-dimensional reconstructions were generated using the Lead-DBS platform^212^.

#### Electrophysiology

Mice were anesthetized with an intraperitoneal injection of 2% Avertin and subsequently perfused with oxygenated (95%O_2_/5%CO_2_) ice-cold cutting solution containing the following: 110mM choline-Cl, 10mM D-glucose, 7mM MgCl2, 2.5mM KCl, 1.25mM NaH_2_PO_4_ • 2H_2_O, 0.5mM CaCl_2_, 1.3mM Na-ascorbate, and 25mM NaHCO_3_. Whole brains were extracted into ice-cold cutting solution and coronal 290 µm brain slices were cut with a vibrating microtome and incubated for 30 minutes in 34°C oxygenated (95%O_2_/5%CO_2_) artificial cerebral spinal fluid (aCSF) with the following: 125mM NaCl, 25mM NaHCO_3_, 2.5mM KCl, 1.25mM NaH_2_PO_4_, 2.0mM CaCl_2_, 1.0mM MgCl_2_, and 25mM D-glucose. ACSF is adjusted to 290-295mOSM by the addition of about 4% H_2_0 by volume. The slices were then stored in oxygenated aCSF at room temperature prior to recording. All recordings were performed at ∼37°C using TC-324B (Warner Instruments). Whole-cell current clamp recordings and intrinsic property measurement were achieved using patch electrodes (tip resistance was 4-6 MOhms) that were filled with a potassium-gluconate internal solution containing the following: 115mM K-gluconate, 10mM HEPES, 2mM EGTA, 20mM KCl, 2mM MgATP, 10mM Naphosphocreatine, and 0.3mM Na_3_GTP. Recordings were sampled at 80 kHz (Multiclamp 700B; Molecular Devices). Current clamp experiments used pipette capacitance neutralization of 3 to 4 pF auto bridge balance adjustments typically between 15 to 35MΩ. Action potentials were captured using template matching to an average action potential with a 1ms baseline and a 4ms length with a minimum separation of 2ms a captured baseline of 10ms with a 40ms length and a threshold of 1. Intrinsic cellular properties were measured using a 10mV test pulse. Spontaneous mEPSCs were detected by recording in 1µM TTX and 10µM SR95531 using template matching in Axograph X. Template parameters included a rise time of 1ms, decay time of 2ms, baseline of 10ms, length 30ms, and had a threshold of 3. mEPSC events were processed with a filter at 2Hz and all captured events were manually reviewed.

#### Surgical preparation and mesoscopic imaging of neuronal activity

Whole-brain expression of GCaMP6s in neurons was achieved by crossing Nkx2.1-Cre; Pten^fl/+^ mice with mice expressing Snap25-GCaMP6s^205^, provided by Hongkui Zeng (Allen Brain Institute). Surgeries and imaging procedures for widefield calcium imaging were as described previously^213^ with modifications described below. Mice were anesthetized using 1 – 2% inhaled isoflurane and were maintained on a recirculating water heating pad for the duration of the surgery. Carprofen (5 mg/kg) and 2% topical lidocaine were administered for analgesia. The skin and fascia layers above the skull were removed to expose the entire dorsal surface of the skull from the posterior edge of the nasal bone to the middle of the interparietal bone, and laterally to the temporal muscles. The skull was thoroughly cleaned with saline and care was taken to not let the skull dry out. The edges of the skin incision were secured using cyanoacrylate (Vetbond, 3M). Small screws (0-80, 3/16”) were secured to the interparietal bone and the nasal bone with Vetbond and transparent dental cement (Metabond, Parkell), and then a thin layer of dental cement was applied to the entire exposed skull. Once the dental cement dried, it was covered with a thin layer cyanoacrylate (Maxi-Cure, Bob Smith Industries). The combination of the dental cement and cyanoacrylate substantially increases the transparency of the skull.

Mice recovered off anesthesia for at least 2 hours prior to imaging. They were then secured by the skull screws to a custom imaging stage with recirculating water heating pad for the duration of imaging. Imaging was performed using a Zeiss Axiozoom V.16 coupled to a PlanNeoFluar Z 1x, 0.25 NA objective with a 56 mm working distance. Epifluorescence excitation is performed using a 7-channel LED driver (SpectraX, Lumencor) mated to the mesoscope by a 3 meter liquid light guide (Lumencor). The blue LED channel is filtered using a ET470/20x filter (Chroma) and the violet LED channel is filtered using a ET395/25x filter (Chroma). LED illumination is reflected onto the imaging plane using a FT495 (HE) dichroic mirror and epifluorescence emissions are filtered with a BP525/50 filter (38 HE, Zeiss) and recorded using a sCMOS camera (Hamamatsu Orca Fusion) with 576×576 resolution after 4×4 pixel binning. Images are acquired by a computer running HCImage software (Hamamatsu). The intensity of the LED excitation at the imaging plane is ∼0.1 mW/mm^2^. Both spontaneous and whisker stimulation-evoked activity was recorded during experiments. For stimulation trials, the mouse’s right whiskers were deflected (a single 100 millisecond deflection every 10 seconds) using a piezo bender (PL112, Physik Instrumente).

#### Preprocecssing of widefield mesoscopic calcium imaging data

Calcium data were preprocessed using BIS-MID^214^. In short, the data were motion corrected, smoothed, down sampled, corrected for photo bleaching, and had a physiological estimate regressed out. Following this the data were moved to a common space, which entailed applying affine transforms to align each scan across subjects. In common space global signal regression (GSR) and band pass filtering were applied. A 2D projected map of the Allen atlas was also transformed into this common space and was used to define seed regions.

#### Rapamycin administration

As previously described^45^, mice received daily intraperitioneal (IP) injections of either rapamycin (10 mg/kg of body weight; Cayman Chemical) or Vehicle Control (5% Tween80 (Fischer Biosciences), and 5% PEG400 (Sigma Aldrich) from P10-P25 for assessment of survival.

#### Everolimus administration

Mice received subcutaneous injections of either everolimus (1 mg/kg of body weight; Cayman Chemical) or Vehicle Control ((5% Tween80 (Fischer Biosciences) and 5% PEG400 (Sigma Aldrich) every 48 hours from P1-P14. The mice were perfused the day after final rapamycin/vehicle treatment at P14 and processed for ventriculomegaly or immunofluorescent labeling analysis as described above.

