## Supplementary Figures for "Dual impact of PTEN mutation on CSF dynamics and cortical networks via the dysregulation of neural precursors and their interneuron descendants"

**A**

| Proband ID | Sex | Ethnicity | Consanguineous | Genotype | Mutation Type | Class | Position (hg19) | AA Change | GnomAD Exome AFR | GnomAD Ge | Bravo Freq. | MetaSVM | MPC | CADD |
|---|---|---|---|---|---|---|---|---|---|---|---|---|---|---|
| KCHYD420-1 | M | European | 0 | Heterozygous | *De Novo* | Stop-gain | 10:89624274–T:G | p.Y16X | 0 | 0 | 0 | . | . | 35 |
| KCHYD487-1 | F | European | 0 | Heterozygous | Transmitted | D-mis | 10:89692905–G:A | p.R130Q | 0 | 0 | 0 | D | 2.50374879 | 33 |
| KCHYD71-1 | M | European | 0 | Heterozygous | *De Novo* | D-mis | 10:89692905–G:A | p.R130Q | 0 | 0 | 0 | D | 2.50374879 | 33 |
| KCHYD381-1 | F | European | 0 | Heterozygous | Transmitted | D-mis | 10:89720763–G:A | :p.S305N | 0 | 0 | 7.96E-06 | D | 1.09590049 | 17.18 |
| KCHYD460-1 | F | European | 0 | Heterozygous | *De Novo* | Stop-gain | 10:89720852–C:T | p.R335X | 0 | 0 | 1.59E-05 | . | . | 40 |

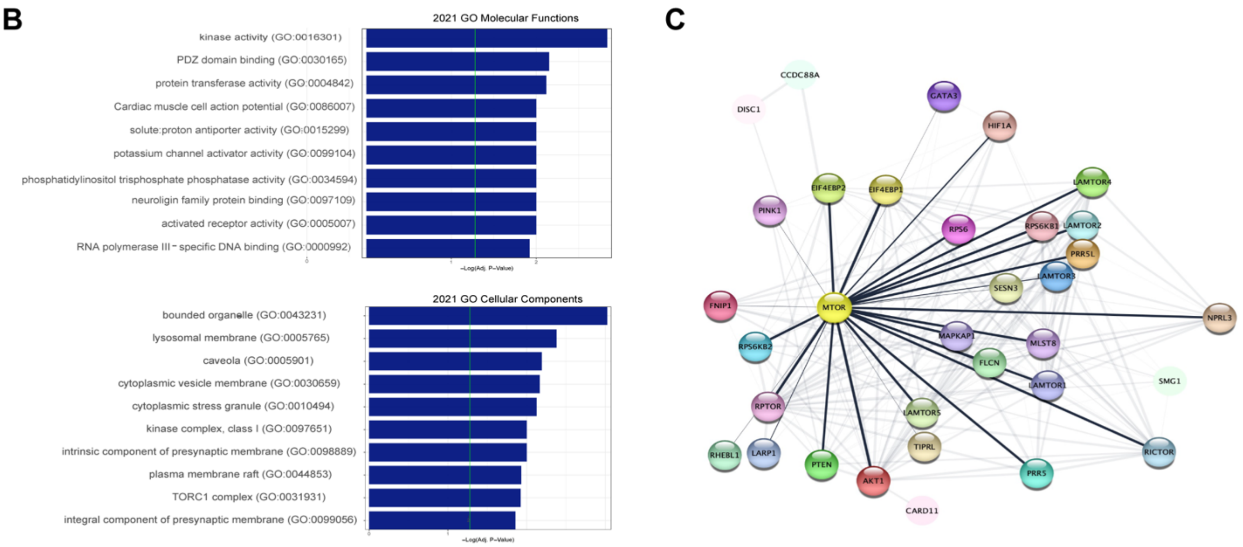


**Figure S1.** (A) PTEN mutations identified in probands from the neurosurgical cohort. (B) 2021 Gene Ontology Molecular Functions and 2021 Gene Ontology Cellular Components from analyses of possible congenital hydrocephalus risk gene list suggest involvement of the PI3K-AKT-mTOR pathway in congenital hydrocephalus pathogenesis. (C) Protein-protein interaction network map of PTEN-mTOR displays a highly-interconnected regulatory network.

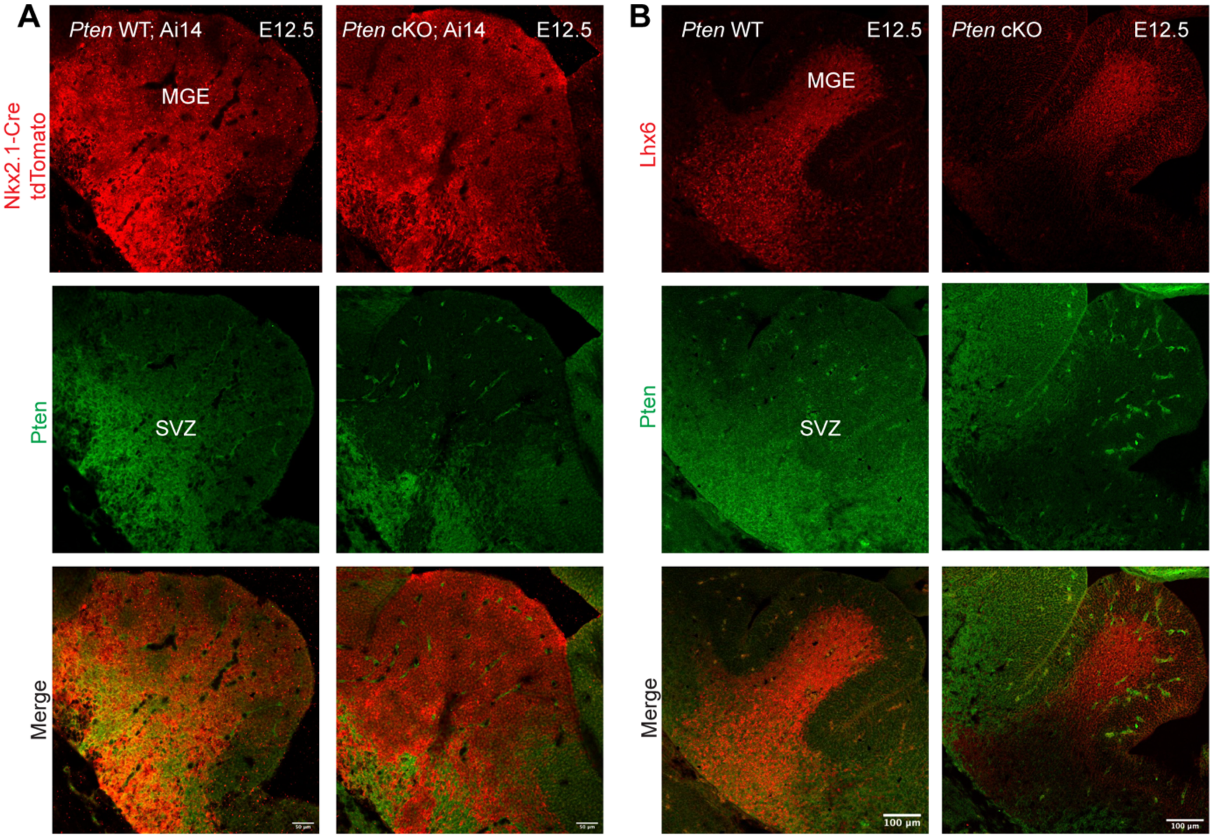


**Figure S2. *Pten* cKO depletes Pten in the Nkx2.1-Cre-expressing NPC lineage within the SVZ of the MGE in mice at E12.5** (A) Immunofluorescent images of E12.5 coronal brain sections showing abundant Pten protein expression in the SVZ of the MGE, defined by Nkx2.1-Cre expression (revealed by the Cre-dependent tdTomato reporter) in *Pten* WT; Ai14+ mice. *Pten* cKO depletes Pte*n* protein expression in the Nkx2.1-Cre expression domains of the MGE. (B) Immunofluorescent images show Pten protein in Nkx2.1-lineage NPCs expressing Lhx6 within the SVZ of the MGE in *Pten* WT mice, and is eliminated in *Pten* cKO mice at E12.5. Scale bar in A =50 μm., Scale bar in B = 100 μm.

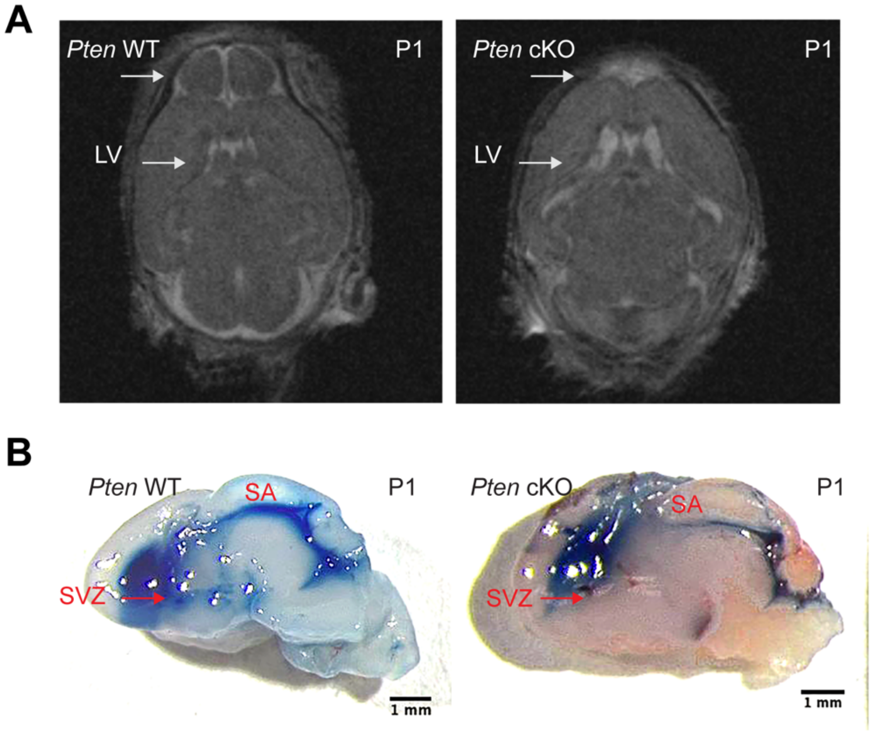


**Figure S3. *Pten* cKO mice display perinatal obstructive hydrocephalus.** (A) Representative Brain MRI at P1 reveals increased size of the lateral ventricles (LV) in *Pten* cKO mutant mice as compared to a littermate control. (B) Representative midsagittal images of dissected brains from a *Pten* WT and *Pten* cKO at P1 ten minutes after ICV injection of a CSF tracer (1% Evan's Blue Dye). White arrowheads highlight decreased olfactory bulbs and enlarged lateral ventricles. Red arrows highlight obstructed CSF flow from the LV, due to prenatal overgrowth of the SVZ in *Pten* cKO mice, causing decreased CSF tracer in the v3V and cerebral aqueduct.

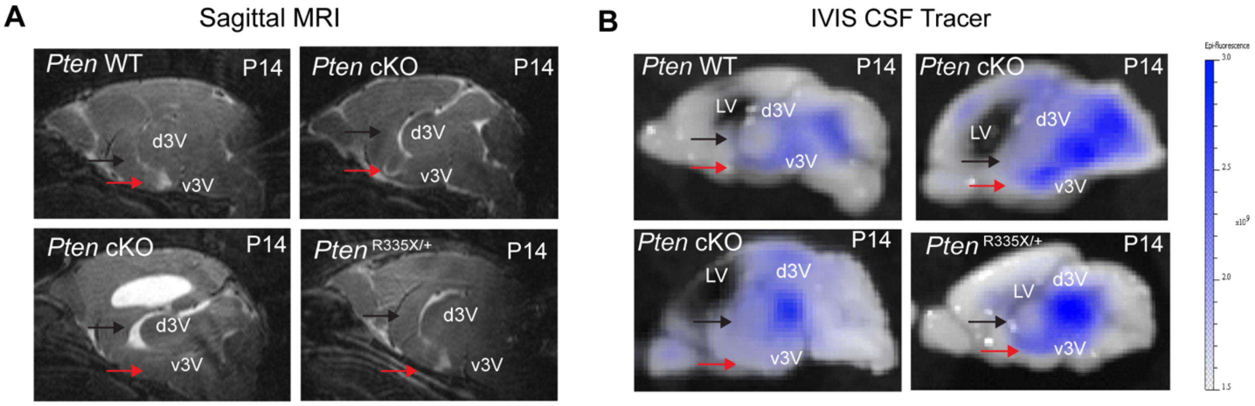


**Figure S4. CSF obstruction does not always occur in ventriculomegalic *Pten* mutant mice**. (A) Representative Brain MRI at P14 reveal increased size of dorsal third ventricle (d3V) of the ventricular systems in *Pten*$^{R335X/+}$ and *Pten* cKO mutant mice. White arrows highlight d3V in *Pten* mutant mice and WT control. Red arrows highlight ventral third ventricle (v3v) caliber in *Pten* cKO mice and WT control. (B) Representative midsagittal images of dissected P14 brains from *Pten* WT and *Pten* cKO mice ten minutes post-ICV injection of CSF tracer Evan's Blue(1%). Black arrows highlight CSF tracer in d3V of *Pten* mutant mice. Red arrows highlight CSF tracer in v3V of *Pten* mutant mice.

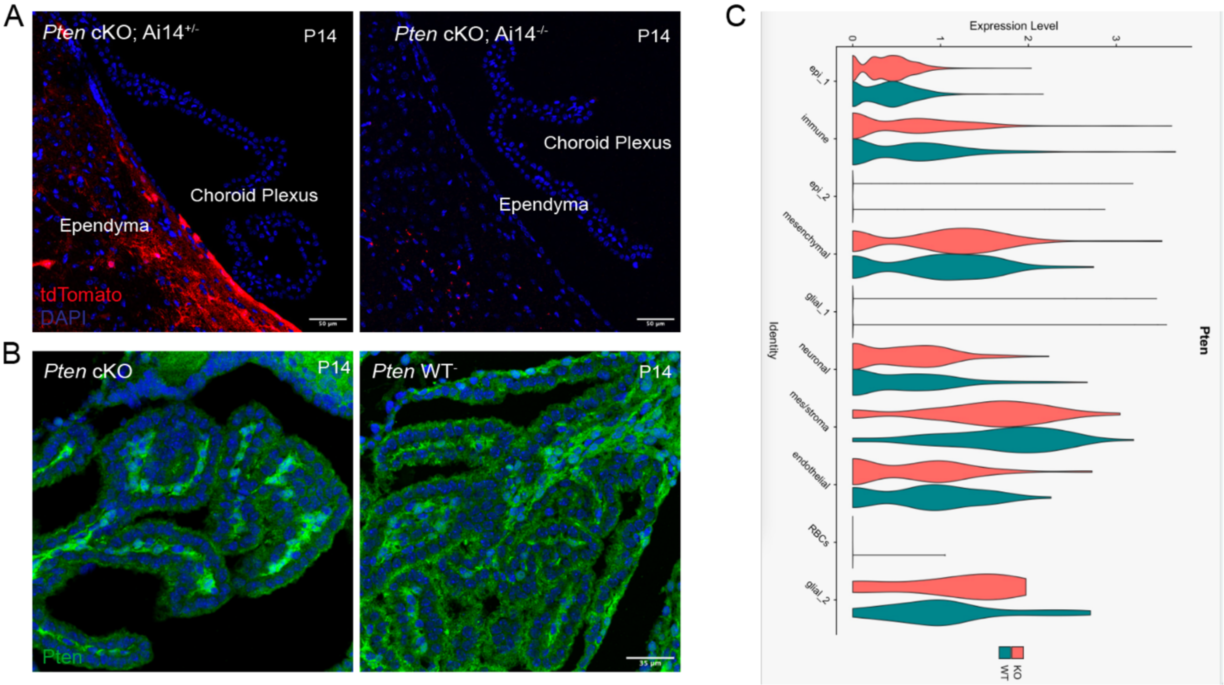


**Figure S5. *Pten* is expressed in the choroid plexus of *Pten* cKO mice**. (A) Representative images of the ChP from a *Pten* cKO mouse with tdTomoto expression (left) and a *Pten* cKO mouse without tdTomato expression (left). Nkx2.1-Cre recombindation occurs within cells in the ependyma and VZ/SVZ (red); however, recombination in the choroid plexus was not detected by expression of the Cre-dependent tdTomato reporter. Scale bars, 50 µm. (B) Representative images of immunofluorescent labelling of the choroid plexus from *Pten* cKO and Pten WT mice reveals preseved Pten protein expression in the choroid plexus in *Pten* cKO at P14. Scale bars, 35 µm. (C) Violin-plots from scRNAseq analysis of *Pten* expression in choroid plexus cells from *Pten* cKO and *Pten* WT mice shows equivalent *Pten* expression between *Pten* cKO and WT controls in each cell cluster.

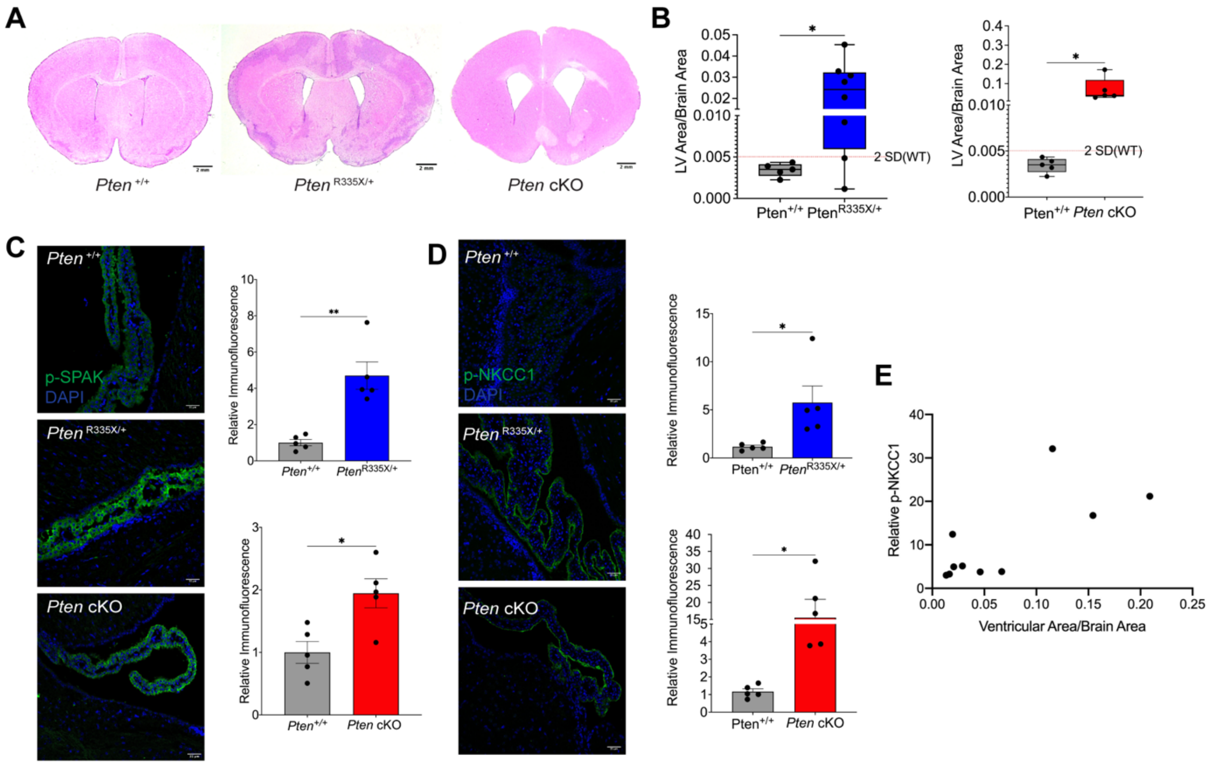


**Figure S6. *Pten* mutant mice have activated SPAK-NKCC1 at the choroid plexus.** (A) Representative images of hematoxalin and eosin staining of coronal brain sections from a *Pten*$^{+/+}$, *Pten*$^{R335X/+}$, and *Pten* cKO mouse at P14 with quantitation in (B) of LV area, normalized by brain area, shows ventriculomegaly in *Pten* mutant mice compared to the same group of control *Pten*$^{+/+}$ mice. Ventriculomegaly defined as LV area/Brain area ratios greater than 2 standard deviations greater than the WT mean (2SD(WT)). (C) Representative images of immunofluorescent labelling of p-SPAK (green) with quantitation reveals significantly increased SPAK activation in the ChP of Pten mutant mice relative to controls (The same group of controls were used for the unpaired, two-tailed t-tests. (D) Representative images of immunofluorescent labelling of p-NKCC1 (green) with quantitation demonstrates significantly increased NKCC1 activation in the ChP of *Pten* mutant mice relative to controls. The same group of controls were used for the unpaired, two-tailed t-tests. (E) *Pten* mutant mice have significant correlation of ratios of LV area/ Brain area with relative p-NKCC1 intensity at the choroid plexus, $p<.05$, significance determined by two tailed, Pearson's test. Error bars, mean ± sem; each symbol represents one animal. *$p<0.05$, **$p<0.01$, ***$p<0.001$, ****$p<0.0001$, ns = not significant.
